# Epstein Barr virus infection induces tissue-resident memory T cells in mucosal lymphoid tissues

**DOI:** 10.1101/2023.11.24.565960

**Authors:** Daniel Kirchmeier, Yun Deng, Lisa Rieble, Fabienne Läderach, Patrick Schuhmachers, Alma Delia Valencia-Camargo, Anita Murer, Nicole Caduff, Bithi Chatterjee, Obinna Chijioke, Kyra Zens, Christian Münz

## Abstract

Epstein Barr virus (EBV) contributes to around 2% of all tumors worldwide. Simultaneously, more than 90% of healthy human adults persistently carry EBV without clinical symptoms. In most EBV carriers it is thought that virus-induced tumorigenesis is prevented by cell-mediated immunity. Specifically, memory CD8^+^ T cells recognize EBV-infected cells during latent and lytic infection.

Using a symptomatic primary infection model, similar to infectious mononucleosis (IM), we found EBV induced CD8^+^ tissue resident memory T cells (TRMs) in mice with a humanized immune system. These human TRMs were preferentially established after intranasal EBV infection in nasal-associated lymphoid tissues (NALT), equivalent to tonsils, the primary site of EBV infection in humans. They expressed canonical TRM markers, including CD69, CD103 and BLIMP-1, as well as Granzyme B, CD107a and CCL5, while demonstrating reduced CD27 expression and proliferation by Ki-67 expression. Despite cytotoxic activity and cytokine production *ex vivo*, these TRMs failed to control EBV viral loads in the NALT during infection although effector memory T cells (TEMs) controlled viral titers in spleen and blood.

Overall, TRMs in mucosal lymphoid tissues are established by EBV infection, but primarily systemic CD8^+^ T cell expansion seems to attenuate viral loads in the context of IM-like infection.

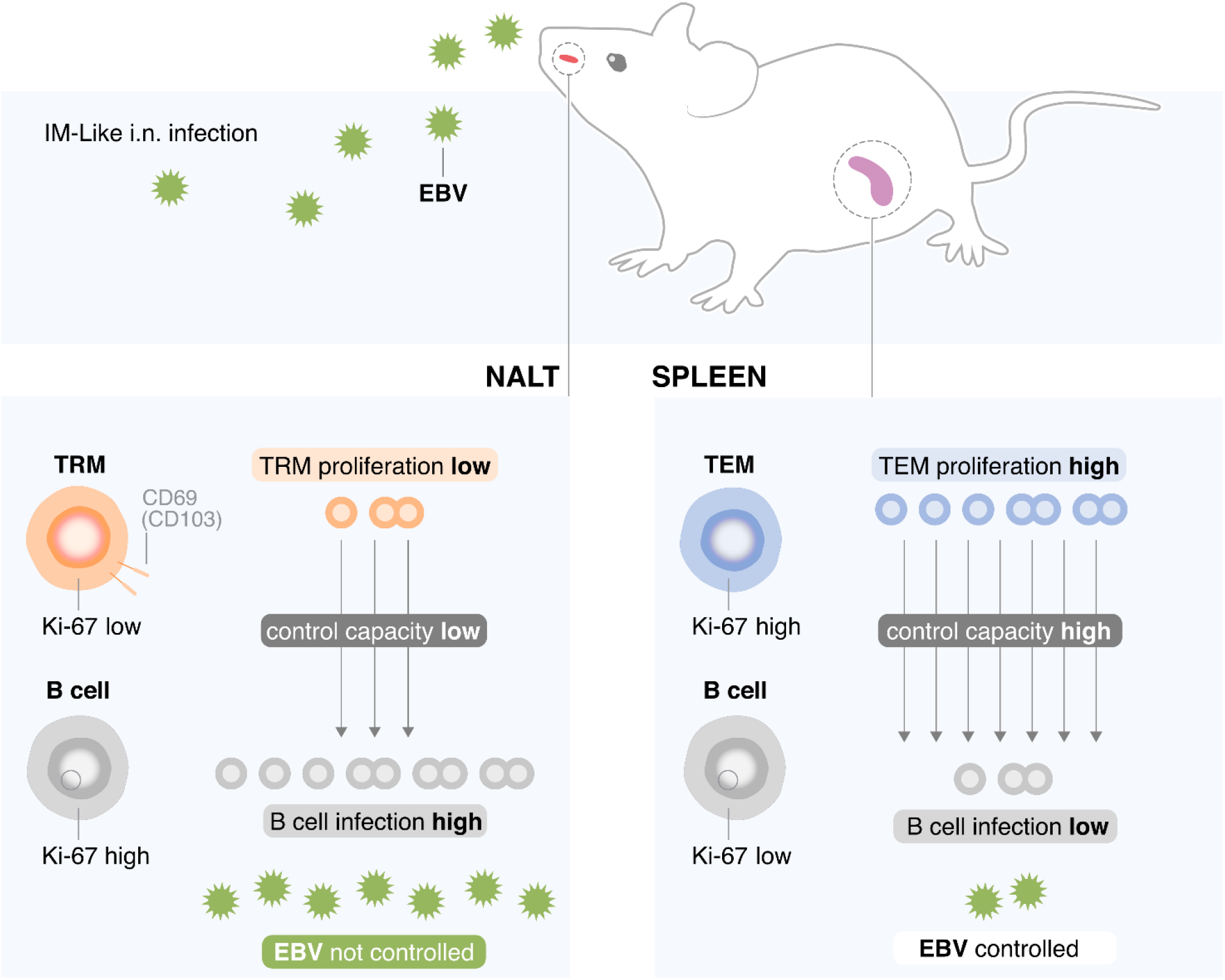

## Introduction

Epstein Barr virus (EBV) or human herpesvirus 4 (HHV4) is a WHO class I carcinogen and associated with approximately 2% of all tumors in humans (1). Although more than 90% of the human adult population is persistently infected with EBV, fortunately only a small number of virus carriers is affected by the around 300,000 new EBV-associated malignancies that arise every year (2–4). EBV is usually transmitted via saliva exchange and infects its main host, the human B cell, in mucosal secondary lymphoid tissues, such as the tonsils, presumably after transcytosis across mucosal epithelia (5, 6). Initially, EBV expresses only latent gene products in B cells that are thought to drive host cell differentiation into the memory B cell pool (7). These EBV nuclear antigens (EBNAs) and latent membrane proteins (LMPs) are then switched off in quiescent memory B cells, which then function as the cellular compartment of long-term viral persistence. Successive antigen-driven B cell differentiation into plasma cells induces lytic EBV replication (8, 9). Although most primary EBV infections are asymptomatic, delayed first encounter with the virus in adolescence or young adulthood increases the risk of developing infectious mononucleosis (IM), a massive CD8^+^ T cell lymphocytosis that is mainly driven by lytic EBV antigens (10, 11). IM usually peaks six weeks after virus encounter (12). Although IM resolves to a persistent, immune-controlled EBV infection in most affected individuals, it nevertheless increases the risk of developing both virus-associated classic Hodgkin’s lymphoma, one of the EBV-associated malignancies, and the autoimmune disease multiple sclerosis (MS) (13, 14). It remains unclear why delayed primary EBV infection increases the risk for IM. High infectious dose or poor and delayed immune control at the initial site of virus encounter, namely mucosal secondary lymphoid tissues such as tonsils, might allow replication to high viral loads. Subsequently this could trigger the systemic CD8^+^ T cell lymphocytosis that causes IM. In order to develop potential therapeutic strategies to alleviate IM, it is important to better understand EBV-specific immune responses during primary infection at mucosal surfaces.

Tissue-resident memory T cells (TRMs) comprise a distinct subset of memory T cells residing stably within tissues. They are best described in the context of pathogens causing self-limiting, acute illnesses and are thought to play central roles in protection from reinfection (15–17). To this extent, a substantial fraction of memory T cells in mucosal tissues are tissue-resident and located at major sites of pathogen encounter. Mucosal TRMs are typically distinguished by expression of CD69, and often co-express the epithelial adhesion molecule CD103 (18–21). While TRMs are generally considered to be important mediators of protective immunity, aberrant TRM generation has been associated with certain pathologies, including asthma, psoriasis, vitiligo, Crohn’s disease and ulcerative colitis (22–26). Furthermore, reduced TRM establishment and function are associated with poor patient outcomes in certain solid cancers, including lung and breast cancers (27–30), and may contribute to reduced control of chronic infections such as human immunodeficiency virus (HIV), hepatitis B (HBV), and hepatitis C virus (HCV) (31–34). TRMs are often thought to represent an intermediate state of memory differentiation between central memory T cells (TCMs) and effector memory T cells (TEMs). Inflammatory conditions skew T cell differentiation towards TEM, whereas less inflammatory conditions tend to favor central memory T cells (TCM) primed for circulation and long-term survival (35). Regarding chronic and persisting infections, it is unclear whether high levels of inflammation or continuous and repeated antigen exposures may negatively impact TRM generation or otherwise reduce their functional capacities.

EBV-specific T cells with a TRM phenotype have been previously described in humans (36–38), notably in the tonsils, which may represent a “frontline” site in the control of viral reactivation and prevention of viral shedding. However, examinations of donor tissues only represent a “snapshot” in the infection process, and the localization and kinetics of EBV-specific memory T cell establishment and function remain incompletely characterized. To this extent, humanized mice, which are engrafted with human lymphocytes and are permissive to EBV infection, represent an ideal model to study T cell response dynamics. In addition, as antigen location is known to be important in the establishment of certain TRM subsets, we have further developed an intranasal model of EBV infection to better recapitulate the natural route of EBV exposure in humans and, perhaps also, the sequence of viral trafficking during the initial course of disease.

Using this model, we have observed the generation of TRMs after primary EBV infection preferentially in nasal-associated lymphoid tissue (NALT), the murine tonsil equivalent. Despite phenotypic similarities to previously described TRM populations and functionality *in vitro*, depletion of CD8^+^ effector memory T cells during EBV infection resulted in increased blood and splenic viral loads, while depletion of TRMs did not influence EBV infection in the NALT. Our data suggest that differentiation stage and an associated disadvantage in proliferative capacity and co-stimulation render TRMs less protective against primary EBV infection with IM, a persisting and, in comparison to other infectious diseases, highly inflammatory infection.

## Results

### EBV initiates infection in nasal associated lymphoid tissues (NALT) after intranasal (i.n.) inoculation

Here we used NOD/SCID/IL2-γ_c_^null^ (NSG) mice and NSG mice harboring an HLA-A*02:01 transgene (NSG-A2), both with human immune system reconstitution (humanized), to investigate cell-mediated immune control of EBV infection *in vivo* (39–41). Using these models, our group previously identified protective roles for circulating CD27^+^2B4^+^CD8^+^ T cells in EBV control (42–45). The establishment and role of mucosal CD8^+^ T cells during EBV infection have, however, not been assessed. To evaluate mucosal T cell responses, we modeled salivary EBV infection by intranasal (i.n.) inoculation with the B95-8 strain expressing luciferase to aid in visualization of infection (45, 46). To ensure consistent establishment of NALT (the murine equivalent of tonsils (47)), we first pretreated experimental animals with a single dose of i.n. administered Staphylococcus enterotoxin B (SEB) two weeks prior to infection, which should mimic bacterial colonization of mucosal surfaces driving post-natal NALT development (48, 49). Following SEB treatment, immunohistochemical analysis of NALTs revealed the accumulation of CD3^+^ and partly CD20^+^ lymphocytes within the NALT (Figure S1A-C). Furthermore, the NALT readily showed CD103^+^ cells along with cells expressing Lyve-1 and peripheral node addressin (PNAd) (Figure S1D-F). This is indicative of lymphatic vessel and high endothelial venule-like structures in a mucosal-associated lymphoid tissue environment.

Following EBV infection, luciferase signal was detectable within the first week post-infection and localized initially to the NALT (Figure 1A). From there, we observed viral spread to the submandibular region, including submandibular and cervical lymph nodes and salivary glands (Figure 1A). Systemic dissemination into the torso was visible from week three onward (Figure 1A-B). In contrast, luciferase signal in the head was detected only rarely following i.p. infection and is mostly attributed to signal from the submandibular region (Figure S1H, 2A-B). We further assessed viral loads in the blood and tissues. Viral loads in the blood had similar kinetics during both i.p. and i.n. infection and this was observed in both humanized NSG and NSG-A2 mice (Figure 1C, S2C,E). In the tissues, EBV was detected in all highly perfused organs, including spleen, liver, and lung following both infection routes (Figure 1DE, S2D). Viral loads were, however, significantly increased in submandibular tissues and NALT following i.n. infection, in comparison to i.p. infection (Figure 1F, S2D). Thus, following i.n. infection, EBV appears to colonize secondary lymphoid tissues of the nasal sinuses before systemically disseminating to the blood and spleen.

**Figure 1:**
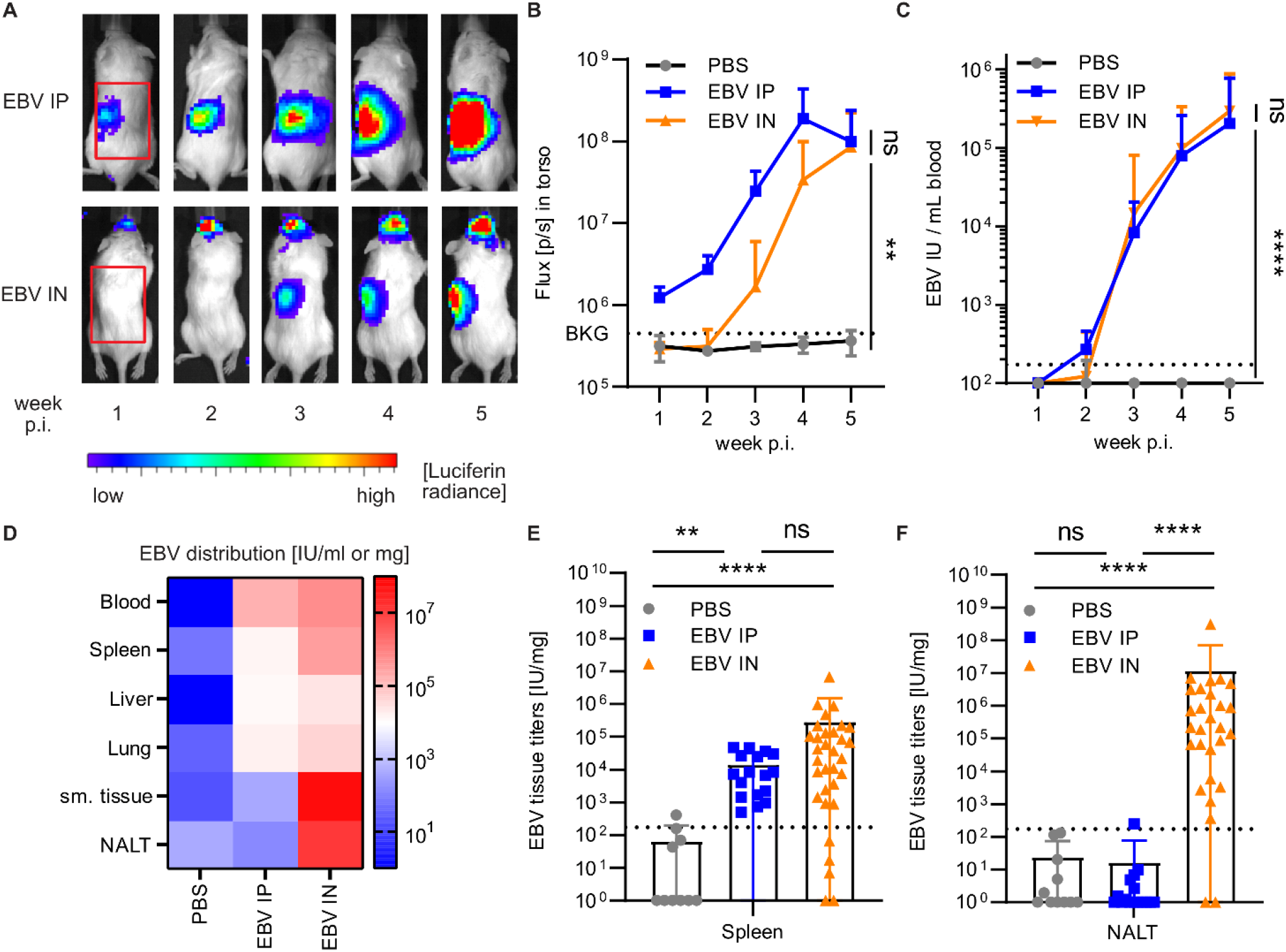
EBV persists additionally in submandibular tissue and NALT upon intranasal infection. (A) Representative IVIS images from week 1 to week 5 after Luc-EBV infection with region of interest (ROI) outlined in red and (B) quantification of the indicated ROI (n=6-24 per group, EBV IP = intraperitoneal EBV infection, EBV IN = intranasal EBV infection, BKG/dotted line = background). (C) EBV viral loads in blood in International Units (IU) / ml of animals over time during EBV infection or PBS controls (n=4-31, see Figure S2C). (D) Heatmap plot for viral titers in indicated tissues or peripheral blood at the termination of the experiment (n = 3-32 per group, see Figure 1E and S2D). (E-F) Quantification of viral loads in spleen or NALT (n=9-33, dotted line at 173 = limit of detection). (B-F pooled data from 3-6 independent experiments (**, p ≤ 0.01; ***, p ≤ 0.001, ****, p ≤ 0.0001, Kruskal-Wallis ANOVA Dunn’s multiple comparison), for EBV IN following animals were used NSG (n=16) and A2-transgenic NSG (n=17)).

### EBV induces both systemic T cell responses and NALT TRMs upon i.n. infection

Route of infection can influence T cell differentiation and subsequent protective capacities (50). We were, therefore, interested in comparing T cell phenotypes following i.p. or i.n. infection. For both infection routes, the CD8^+^ to CD4^+^ T cell ratio in peripheral blood increased progressively over time, indicating CD8^+^ T cell-dominated responses (Figure 2A). Thus, we focused our further analyses on CD8^+^ T cells. Following both i.p. and i.n. infection, CD8^+^ T cells shifted away from a CD45RA^+^CD62L^+^ naïve phenotype (Figure 2B), with concomitant expansion of CD45RA^-^CD62L^+^ central memory cells (TCMs) and CD45RA^-^CD62L^-^ effector memory cells (TEMs) (Figure 2C). For both infection routes, we further observed high frequencies of TEMs in both highly blood perfused tissues, including spleen, liver, and lung, as well as in mucosal tissues, including the NALT (Figure 2D,F, S3A), indicating that activated effectors traffic throughout much of the body during infection. Interestingly, following i.n. EBV infection, both frequencies and numbers of CD69^+^ TEMs and CD69^+^CD103^+^ TEMs were increased in the NALT (Figure 2E, G-H, S3B). These cells were further predominantly intravascular antibody negative (Figure 2I), suggesting that they were TRMs (51). Thus, while both i.p. and i.n. EBV infection elicit systemic TEM responses, i.n. infection selectively expands T cells with a TRM phenotype in the NALT.

**Figure 2:**
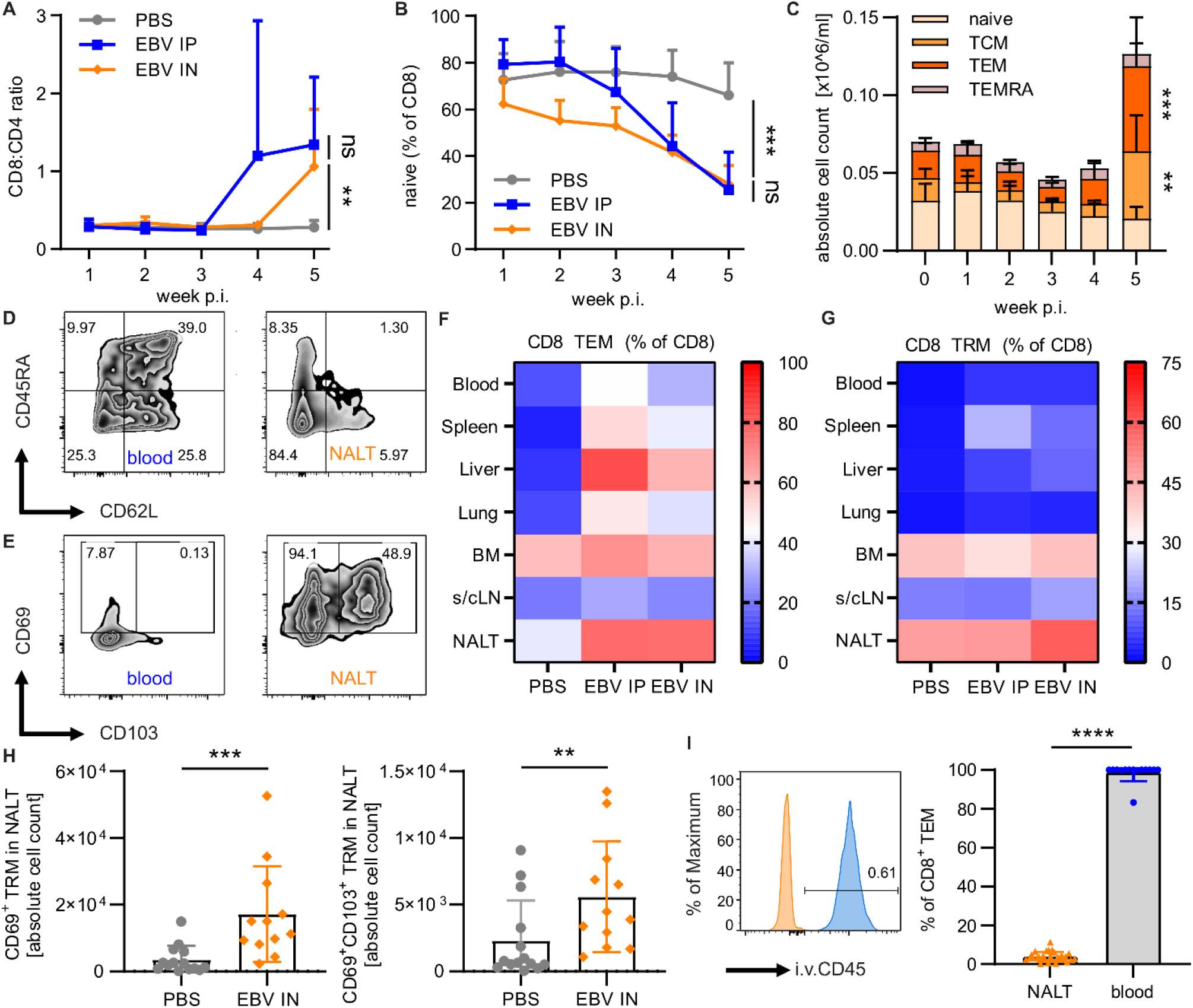
EBV infection induces TRM cells in the NALT. (A) Ratio of CD8:CD4 T cell frequencies and (B) frequency of naïve (CD45RA^+^CD62L^+^) in blood over time measured by flow cytometry. (C) Numbers of naïve, central memory (CD45RA^-^CD62L^+^, TEM), effector memory (CD45RA^-^CD62L^-^, TEM) and TEMRA (CD45RA^+^CD62L^-^) CD8^+^ T cells after EBV intranasal infection (EBV IN) in blood over time measured by flow cytometry. (D) Representative flow cytometry plots and gating of CD45RA against CD62L on CD8^+^ T cells and (E) CD69 against CD103 on TEM CD8^+^ T cells of blood and NALT. (F) Heatmap plot representing frequencies of TEM (CD45RA^-^ CD62L^-^) and (G) TRM (CD69^+^ TEM) of CD8^+^ T cells in multiple investigated organs (n= from 4-34, see Figure S3C-D). (H) Quantification of absolute cell numbers of CD69^+^ TRMs (left) and CD69^+^CD103^+^ TRMs (right) in the NALT in PBS versus intranasally infected mice (EBV IN). (I) Overlay of a representative flow cytometry plot of i.v. injected CD45-AF700 in NALT (orange, n=28) and blood (blue, n=14) and quantification. Pooled data (n=2-9, kinetics depicted as means with 95% CI, *, P ≤ 0.05; **, P ≤ 0.01, ****, P ≤ 0.0001, Kruskal Wallis ANOVA for three groups or Mann Whitney U-test for comparison of two groups)

### Single cell-RNA-Sequencing of NALT T cells reveals TRM transcriptome

To characterize the transcriptional profile of the NALT T cells elicited by i.n. infection, we performed single-cell RNA-sequencing (scRNA-seq) of CD8^+^ TEMs (to enrich for virus-specific cells) sorted from blood, spleen, submandibular and cervical LN (draining the palatine of the nasopharyngeal cavity, head and neck) and the NALT of EBV-infected animals. This allowed us to compare the patterns of transcription between NALT T cells and T cells from organs not harboring a pronounced resident phenotype during infection.

Uniform Manifold Approximation and Projection (UMAP) plots of the scRNA-seq data revealed that NALT cells were transcriptionally distinct from those of the other three organs (Figure 3A). Using an unsupervised clustering algorithm (52), we identified nine distinct clusters within the four tissues, including two clusters within the NALT (Figure 3B and S4A, respectively). We further computed pseudo-temporal cell relationships between the identified clusters using the TSCAN algorithm to evaluate cellular developmental trajectories (53). We found that the two NALT clusters were likely derived from clusters in the center of the UMAP (Figure 3C), suggesting that they differentiated from other cell populations shortly after migrating into the NALT. Within the two NALT clusters, however, patterns of differential gene expression appeared to be comparable (Figure S4B). By differential expression analyses (54), we next identified the 50 genes most up- and down-regulated by NALT cells, compared to those of other tissues (Figure 3D). Genes previously reported to be important for TRM development including *RUNX3*, *BHLHE40,* and *PRDM1* (encoding BLIMP-1), as well as markers described as TRM core signature genes, including PDCD1 (encoding PD-1) and CD69, were upregulated in NALT-derived T cells (55–58). Genes encoding cytokines and cytotoxic molecules, including *IFNG*, *CCL3*/*CCL4*/*CCL5* and *GZMB*, were also elevated in cells from the NALT. When plotting selected markers of interest onto the UMAP, TRM-related gene expression was predominately increased in NALT cells, while *KLF2* and *S1PR1*, associated with tissue egress, were more highly expressed in cells derived from the blood and spleen (Figure 3E) (59). Lastly, NALT-derived cells further displayed increased *CXCR4* and *LAG3* transcription compared to cells from other organs (Figure 3D). Altogether, these data support that NALT-derived T cells responding to EBV infection have a TRM-like transcriptional profile and likely represent a *bona fide* population of TRMs that is generated in this tissue in response to EBV infection.

**Figure 3:**
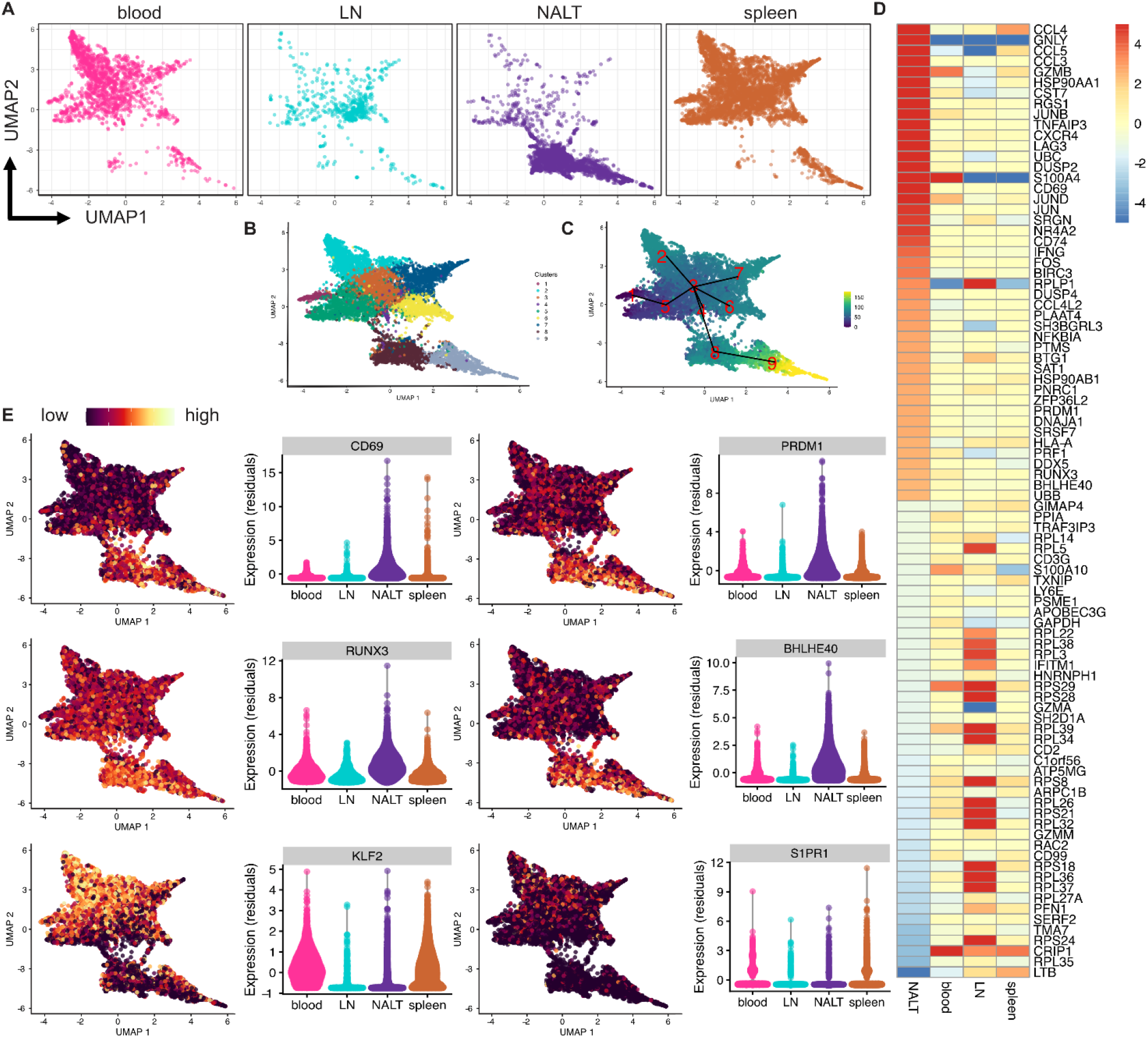
Single cell-RNA-Sequencing reveals TRM transcriptome in the NALT of EBV i.n. infected humanized mice. (A) UMAP plot of single CD8^+^ TEM transcriptomes of EBV infected humanized mice in the respective tissues. (B) Integrated UMAP plot of unsupervised clustering and (C) cellular trajectories in pseudo time (blue to yellow = earlier to later pseudo time point) of TEM transcriptomes of all tissues. (D) Heatmap showing log2(fold change) of top 50 of highest and lowest differentially expressed genes of NALT single cell transcriptomes (rows). (E) Integrated UMAP of cells showing relative expression of chosen TRM and circulating T cell marker expression (yellow = high expression, red = intermediate expression, black = low expression). Next to each plot are violin plots showing individual expression of the respective markers as normalized residual values.

In addition to evaluating polyclonal responses, we utilized an adoptive transfer model to confirm the establishment of EBV-specific CD8^+^ TRMs following infection. To that end, we transduced splenocytes of humanized mice *ex vivo* with HLA-A*02:01 restricted EBV-specific T cell receptors (TCR) recognizing either lytic BMLF1 or latent LMP2 antigens, as described previously (44, 45). These EBV-specific TCR-transgenic cells were then adoptively transferred into autologously reconstituted littermate animals (Figure S5A). Transduced cells were identified by expression of both the respective HLA-A*02:01/peptide pentamer and the murine membrane-proximal TCR domain (due to the hybrid human/mouse format of the transgenic TCRs, Figure S5B). TCR-transgenic cells were able to migrate to the NALT and generated both CD69^+^ and CD69^+^CD103^+^ TRMs (Figure S5C-F). Of note, TCR-transgenic cells were detected in the NALT but at much lower frequencies than in the spleen, which prevented reliable quantification but suggests that EBV-specific T cells undergo lower expansion in the NALT compared to the spleen. Alternatively, migration into the NALT could require a specific phenotype that these TCR-transgenic cells do not easily acquire. To further investigate the relationship between CD8^+^ TCR clonotypes of T cells from NALT and spleen, we analyzed the TCR-Vβ repertoires (Figure S6A). Interestingly, we found that TCR-Vβ chain usage of expanded CD8^+^ T cells clones in the spleen differed substantially between individual EBV-infected mice (Figure S6B). Using the six TCR-Vβ chains detected in the spleen with the highest frequencies in every mouse, we analyzed the overlap with TCR-Vβ chains from the NALT of the same individuals. Intriguingly, within individual mice, we observed that the most expanded TCR-Vβ chains overlapped in both spleen and NALT (Figure S6C). Thus, combined with our pseudo-temporal gene expression analysis, these data are consistent with a model whereby, upon EBV infection, a given virus-specific T cell is clonally expanded and gives rise to daughters which populate both the NALT, where they differentiate into TRM, and the spleen, where they remain predominantly TEM.

### Comparison of TRM markers between NALT of EBV-infected humanized mice and human tonsils

We were additionally interested in how NALT T cells from EBV-infected humanized mice (which are predominantly CD69^+^ TRMs) compare to T cells from healthy human tonsils. Thus, we analyzed samples collected from organ donors or tonsillectomies, investigating their T cell and TRM phenotypes. Compared to NALT from EBV-infected animals, human tonsils exhibited higher frequencies of T effector memory cells re-expressing CD45RA (TEMRAs) (Figure 4A, 2D). However, tonsils also contained substantially increased frequencies of CD69^+^ and CD69^+^CD103^+^ TEMs (TRMs) compared to human blood (Figure 2E, 4B-C). In addition, the expression of TRM-related surface markers such as CD11a and PD-1 was increased in NALT and tonsils compared to TEMs in control tissues including spleen or peripheral blood (Figure 4D-G). Intracellular transcription factors and chemokines only partially overlapped between TEMs of the two organs. In particular, the hallmark TRM transcription factor BLIMP-1 was consistently expressed in the NALT, whereas only a subset of tonsils contained a prominent BLIMP-1^+^ population (Figure S7A-B, E-F). Cytotoxic molecules and chemokines, including granzyme B and CCL5, were prominently expressed in the NALT, but only CCL5 expression was increased in tonsils compared to blood (Figure S7C-D, G-H). Interestingly, in the NALT, expression of the costimulatory molecule CD27, required for efficient immune control of EBV (45), was lower than in the spleen of EBV-infected animals (Figure 4D-E). In the tonsils, CD27 expression was intermediate; lower than that observed for naïve CD8^+^ T cells but higher than that of CD8^+^ TEMs in the blood (Figure 4F-G). Accordingly, CD8^+^ TEMs found in the blood of healthy human donors are primarily specific for other pathogens that drive CD27^-^ memory T cell differentiation (60, 61). For NALT TEMs, the expression of TRM-related surface markers further coincided with a lack of S1PR1 expression (Figure S7I-J). Therefore, primary EBV infection might increase granzyme B and BLIMP-1 in TRMs, but seems to diminish CD27 expression, important for EBV immune control.

**Figure 4:**
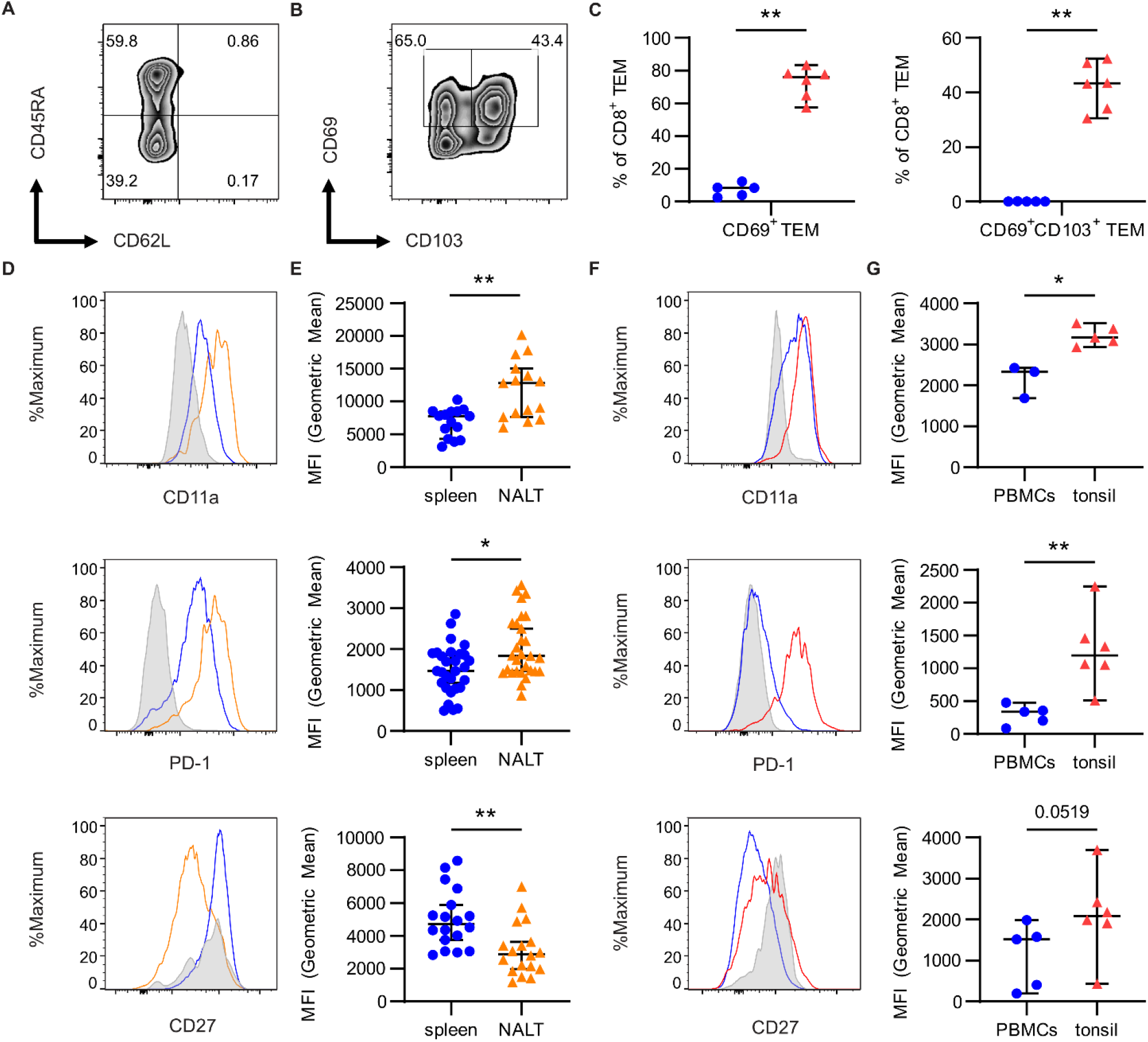
Similarities in TRM markers between NALT TEMs and tonsillar TRMs. (A) Representative flow cytometry gating for identification of CD8^+^ TEM and (B) CD69^+^ and CD69^+^CD103^+^CD8^+^ TEM in human tonsils and (C) quantification in human tonsils (red) and human PBMCs (blue). (D) Histograms and (E) geometric mean fluorescence intensity (MFI) of indicated phenotypic markers in NALT-derived TEMs (orange), TEM from spleen (blue) and naïve CD8^+^ T cells from spleen (gray) of EBV-infected humanized mice. (F) Histograms and (G) MFI of indicated phenotypic markers in CD69^+^CD103^+^ TRMs of human tonsils (red) and TEM of human PBMCs (blue) and naïve (gray) CD8^+^ T cells from PBMCs. To determine significance, Mann-Whitney-U test was performed (*P < .05, **P < .01) and plots represent at least six (humanized mice) or two (human tonsils) independent experiments. For tonsils: n = 5-6 human subjects (except CD11a in tonsil/PBMC is from one experiment comprising n = 3-5 subjects).

### EBV induced NALT TEMs are functional in vitro

We next compared the functional capacity of TEMs derived from the NALT (predominantly CD69^+^ TRMs) and spleen (predominantly CD69^-^ TEMs) of EBV-infected animals or PBS controls, as well as of sorted CD69^-^ and CD69^+^ TEMs (TEM and TRM, respectively) from healthy human tonsils. Following *ex vivo* stimulation, NALT-derived cells from infected animals demonstrated increased IFN-γ and TNF-α co-expression compared to non-infected PBS controls (Figure 5A,C-D top panels). Surface expression of CD107a, as well as granzyme B expression, were also increased, indicating enhanced degranulation and cytolytic capacities (Figure 5B,C-D bottom panels). NALT cells from infected animals further expressed higher levels of granzyme B compared to corresponding spleen TEMs (Figure S7C-D), although cytokine and CD107a expression were similar (data not shown). When comparing CD69^-^ TEMs and CD69^+^ TRMs from human tonsils, we observed only subtle differences (Figure 5E-H). Surface expression of CD107a was pronounced in both NALT (Figure 5B) and tonsillar TRMs (Figure 5F). While overall granzyme B expression by tonsillar TRMs was less pronounced compared to NALT TEMs, there was a trend toward co-expression of CD107a and granzyme B in tonsillar TRMs, which was also observed in NALT TEMs (Figure 5D,H). Unstimulated tonsillar and NALT T cells did not degranulate nor produce cytokines (Figure S8). Together, these data suggest that both NALT and tonsillar TRMs are multifunctional (IFN-γ, TNF-α) and possess the ability to degranulate and express cytolytic molecules.

**Figure 5:**
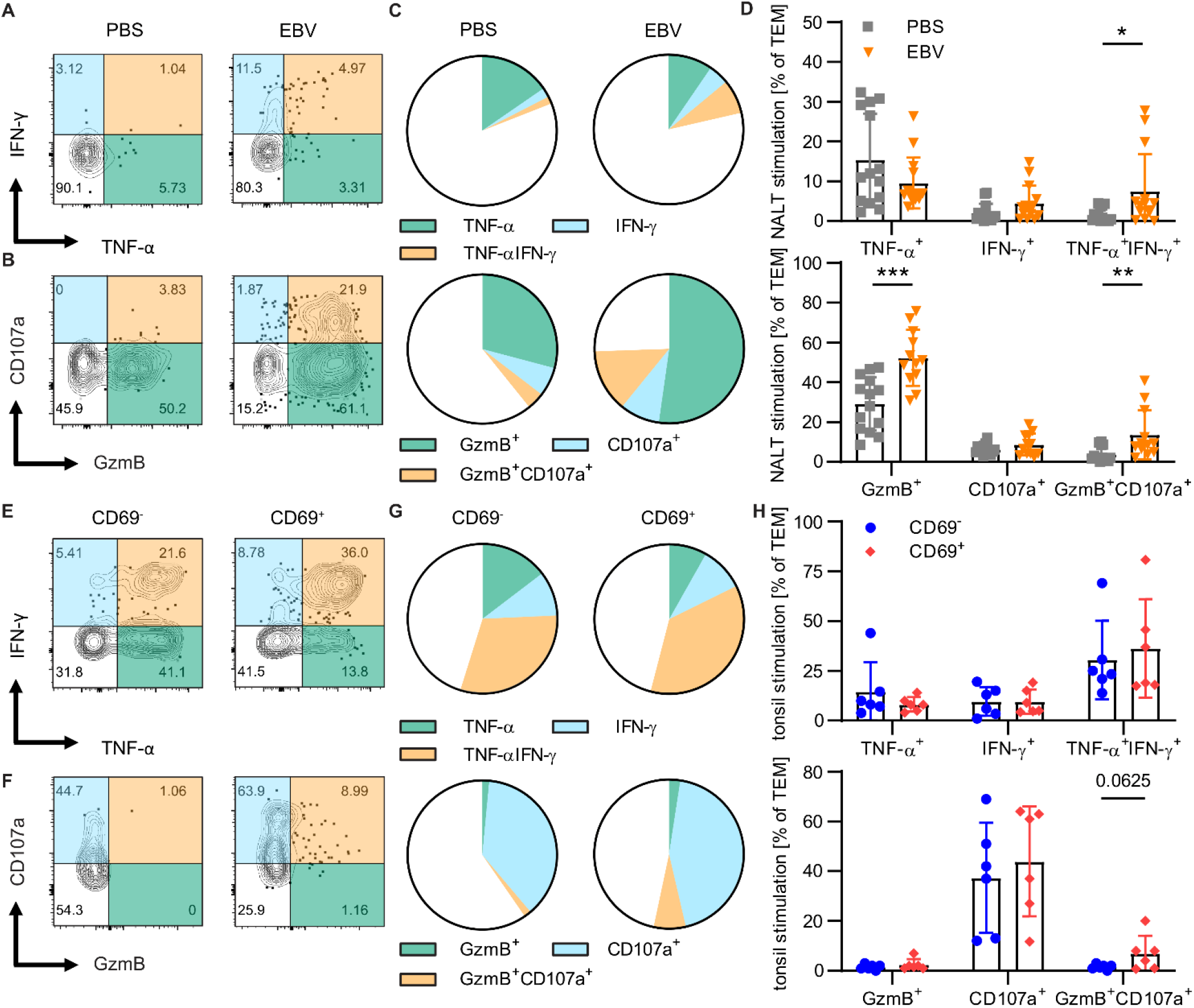
EBV-induced NALT TEM are functional *in vitro*. (A) Representative flow cytometry plots showing IFN-γ and TNF-α or (B) CD107a and granzyme B (GzmB) expression by CD8^+^ TEMs in NALT of non-infected (PBS) or EBV infected individuals after stimulation. (C) Distribution of respective CD8^+^ TEM population and (D) quantification in PBS (gray) and EBV infected (orange) NALT TEMs. (E) Representative flow cytometry plot showing IFN-γ and TNF-α or (F) CD107a and GzmB expression by CD69^-^ or CD69^+^ human tonsil CD8^+^ TEMs. (G) Distribution of respective CD8^+^ TEM population and (H) quantification in CD69^-^ (blue) and CD69^+^ (red) human tonsil CD8^+^ TEMs. (A-C,E-G) Respective quadrants of flow cytometry plots are colored and represented in pie charts (C,G) using the same color. To determine significance Wilcoxon matched-pairs signed rank test (for humanized mouse tissues) or Mann-Whitney U test (for human tonsils) were performed (*P < .05, **P < .01, ***P < .001, data from at least two independent experiments)

### T cell depletion leads to increased EBV viral loads in blood and spleen, but not in the NALT

After characterizing cytokine production and cytolytic potential of NALT TRMs, we next assessed their functional role *in vivo* by antibody-mediated depletion. To allow for accumulation of TRMs in the NALT, we depleted CD8^+^ T cells by OKT-8 antibody injection beginning at week three post-intranasal EBV infection. CD8^+^ T cells in NALT, spleen and blood were significantly reduced in treated animals (Figure 6A). In order to avoid any potential bias due to down-regulation of CD8 expression, we assessed CD4^-^ CD3^+^ T cell frequencies as a measure for CD8^+^ T cell depletion efficacy. The residual CD4^-^CD3^+^ T cell population was more common in spleen and blood than in NALT, suggesting more efficient T cell depletion and a reduced capacity of CD8^+^ T cells to expand in the NALT compared to spleen and blood. Furthermore, this indicated that in contrast to intravascular antibody labelling, which is of a short duration, prolonged treatment with CD8 depleting antibody was able to efficiently label NALT CD8^+^ T cells for depletion. CD8^+^ T cell depletion resulted in a marked increase in viral loads in the peripheral blood during infection (Figure 6B), also indicative of viral spread from the initial site of infection. Strikingly, depletion did not affect viral titers in the NALT (Figure 6C), although viral loads increased in the spleen and blood of depleted animals (Figure 6D-E). To confirm the limited effect of NALT TRMs on EBV control, we specifically targeted the CD103^+^ TRM compartment by treatment with an anti-CD103 antibody (Ber-ACT8) at three weeks post-i.n. EBV infection. We observed a substantial decrease in CD69^+^CD103^+^ double-positive TRMs, although the frequency of CD69^+^ single-positive TRMs did not change (Figure S9A-B). Significantly, viral loads, including titers in the NALT, did not change upon CD103 targeting, confirming the results obtained by the systemic T cell depletion approach (Figure S9C). To decipher further functional differences between lymphocytes within NALT and spleen, we assessed proliferative capacities of T and B cells in both organs. Interestingly, our findings revealed that NALT TEMs exhibit substantially lower expression of the proliferation marker Ki-67 compared to TEMs in the spleen (Figure 6F). In contrast, B cells in the NALT demonstrated significantly higher Ki-67 expression compared to B cells in the spleen (Figure 6G). The observation that CD8^+^ T cells appear to control EBV viral loads in blood and spleen, but not in the NALT, during IM-like infection suggests that TRMs are less potent than TEMs in controlling primary IM-like EBV infection. Together, these findings suggest a model in which NALT TRMs, while functional, are unable to control the robust local B cell proliferation, leading to poor immune control and viral spreading. On the other hand, spleen TEMs, partly due to their superior proliferative capacity, are able to better control the expansion of virally infected B cells.

**Figure 6:**
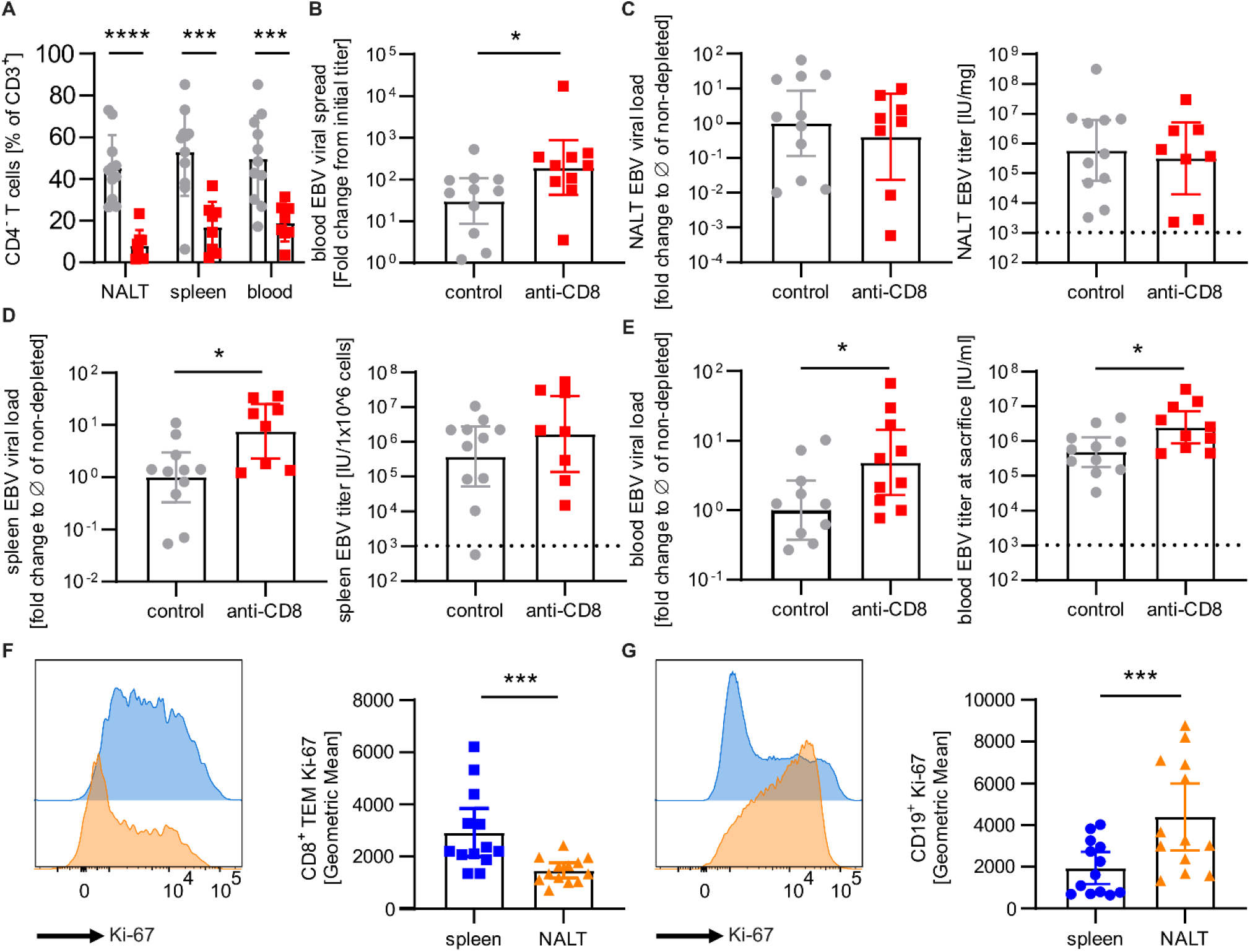
Upon depletion of CD8^+^ T cells, EBV viral loads increased in spleen and blood, but not in the NALT. (A) CD8^+^ cell depletion efficacy analysis by flow cytometry in NALT, spleen and blood of non-depleted control (gray) and anti-CD8 depleted (red) HLA-A2*02:01 transgenic humanized NSG mice as % of CD4^-^ CD3^+^ T cells. (B) EBV viral spread in blood (fold change from initial viral burden to sacrifice). (C) EBV viral loads in International Units (IU) / mg NALT tissue (left) and the matching relative viral loads normalized to the mean of the corresponding nondepleted group (right). (D) EBV viral loads in International Units (IU) / 1×10^6 splenocytes (left) and the matching relative viral loads normalized to the mean of the corresponding nondepleted group (right). (E) EBV viral loads in International Units (IU) / ml blood at sacrifice and the matching relative viral loads normalized to the mean of the corre sponding nondepleted group (right). (F) Representative flow cytometry plot showing Ki-67 expression of CD8^+^ TEM cells in spleen (blue) and NALT (orange) and the respective quantification. (G) Representative flow cytometry plot showing Ki-67 expression of CD19^+^ cells in spleen (blue) and NALT (orange) and the respective quantification. Pooled data from two independent experiments (*, P ≤ 0.05; ***, P ≤ 0.001, ****, P ≤ 0.0001, A-E: Mann Whitney U-test, F-G: Wilcoxon matched-pairs signed rank test, dotted line = limit of detection).

## Discussion

In this study we identified a CD8^+^ TRM population that arises in the NALT after intranasal, but not intraperitoneal EBV infection of humanized mice. This underscores the importance of initial antigen location for TRM generation in mucosal lymphoid tissues (50, 62). These TRMs accumulate during EBV infection and display phenotypic and functional characteristics similar to TRMs of other tissues. However, in comparison to early differentiated CD27^+^ effector memory T (TEM) cells that massively expand in lymph nodes and spleen during EBV infection, NALT TRMs seem to have limited capacity to restrict EBV infection in submucosal secondary lymphoid tissues. EBV might have adapted to initial infection at submucosal secondary lymphoid tissues that allow for frequent re-infections (63, 64).

Even prior to infection, the NALT contains a low number of TRMs. This indicates its potential function as a niche for mucosal TRMs, similar to other lymphoid tissues adjacent to body barriers or long-term niches such as the bone marrow (65–67). Following EBV infection, we detected an increase in the TRM population. This is in line with previous studies demonstrating TRM expansion in the NALT after Influenza infection (68, 69). It was additionally shown that EBV-specific T cells can comprise up to 20% of all T cells within the tonsils (36). This T cell accumulation might be driven by EBV-induced B cell proliferation during IM-like primary infection (70, 71). EBV-infected B cells within the NALT seem to expand similarly during infection, which likely leads to enhanced T cell recruitment and stimulation in the NALT.

The NALT is considered an inductive site for humoral and cellular immune responses in the upper respiratory tract (47, 72). Following respiratory virus infections, CD103^+^ T cells are induced in the NALT as well as throughout the nasal turbinate and septum (66). Human studies have shown that CD103^+^ TRMs locate within the tonsillar lymphoepithelial barrier, indicating a patrolling function towards the epithelium (37). The retention of TRMs within the NALT is likely achieved by expression of CD69, CD103 and other integrins including CD11a, which is known to be associated with the retention of memory T cells in other mucosal tissues, such as the lung (73). Compared to previously described TRMs, we could not confirm any specific chemokine receptor expressed by NALT TRMs. They did not express CXCR6, which was previously defined for TRM homing to lung (74). While a subset of TRMs in the NALT expressed *CXCR4* transcripts, the respective protein was not detected by flow cytometry (data not shown). However, in a skin model it was shown that CXCR4 is dispensable for TRM migration (75). The transcriptional profile of EBV-induced NALT TRMs is consistent with previously described TRM populations, including the expression of TRM core transcription factors *PRDM1* (BLIMP-1), *RUNX3* and *BHLHE40*. BLIMP-1 and RUNX3 were previously shown to be indispensable for TRM development and function, while BHLHE40 was identified as master regulator for TRM metabolism (55, 56, 58). Thus, the NALT appears to serve as a niche that allows the development of *bona fide* TRMs.

In the setting of IM-like disease in humanized mice, NALT-localized T cells did not appear to be essential for local viral control. We recently reported that CD27^+^ T cells are protective against EBV (45), and CD27-deficiency, as well as loss of its ligand, CD70, in humans is associated with persisting EBV viremia and associated pathology (76, 77). Recently, CD27 was additionally described to be one of five markers predictive for EBV-specific T cells within human blood samples (61). Intriguingly, we observed that NALT T cells expressed lower levels of CD27 than their counterparts in the blood and spleen, potentially explaining their reduced EBV-specific immune control in this setting. Similarly, it has been suggested that TRMs are less efficient in the immune control of EBV-infected B cells in the central nervous system (CNS) of MS patients, but might contribute to the inflammatory environment in the CNS during this autoimmune disease (78).

Efficient TRM generation and function seem to depend on conditions of intermediate inflammation. Indeed, reduced T-bet expression among effectors has been shown to favor TRM, as opposed to TEM, establishment (79). As IM represents a more inflammatory manifestation of primary EBV infection, it is possible that this environment favors the establishment of TEM or TEMRA-like T cell memory. Indeed, we observed enhanced TRM generation in a low-dose infection model where viral replication is more moderate and T cell expansion more gradual (44). Similarly, in the context of IM-like EBV infection, T cells express inhibitory molecules including PD-1 and LAG-3, which could potentially be used by EBV-infected B cells to inhibit their function. Interestingly, previous work has demonstrated that, in the tonsils of individuals with IM, fewer memory T cells, including TRMs, are established compared to the tonsils of asymptomatically infected viral carriers (36). These findings are consistent with a scenario where more inflammatory disease prevents efficient establishment of TRMs. It would be interesting to assess whether this further correlates with later indicators of poor EBV immune control such as prolonged viral shedding in the throat, but not blood, of IM patients (80), or increased incidence of EBV-associated malignancies. Perhaps efficient TRM establishment is important in the later, long-term control of such EBV reactivation-associated conditions.

That the EBV-elicited TRMs observed in this study lacked expression of co-stimulatory molecules crucial for EBV-specific immune control could explain the unchanged viral titers upon T cell or CD103^+^ TRM depletion in the NALT. Notably, our model may not reveal protective TRM functions later during persistent EBV infection; EBV infections in humanized mice were only possible for up to six weeks, at which timepoint viral loads peak in IM patients and plateau in humanized mice (12, 45, 81). Due to immune pathology in response to this high viral load, EBV-infected humanized mice needed to be euthanized at this timepoint. This prevented assessment of TRM phenotype and function at later timepoints when a lower setpoint of persistent viral loads is reached in most IM patients. However, after six weeks of infection with plateauing EBV loads, the NALT nearly uniformly contained T cells with a tissue-resident phenotype. In contrast, in tonsils from presumably healthy virus carriers, a mixture of T cell subsets was observed but was enriched for TRMs compared to spleens of humanized mice after six weeks of EBV infection. Notably, this mixture contains CD8^+^ T cells with higher proliferative potential expressing co-stimulatory molecules of early differentiated CD8^+^ T cells, including CD27, that are required for EBV-specific immune control (45). In contrast, in this context, more terminally differentiated TRMs, predominating in the NALT with reduced proliferative capacities (82), do not seem to be beneficial in controlling EBV infection and associated pathology, despite similar clonal TCR-Vβ expansions compared to spleens of the same animals. This could explain the diminished immune control of EBV in the mucosa during IM, resulting in prolonged viral shedding into the saliva for months compared to control of viral loads in the blood within weeks after peak viremia (80). Taken together, our results, along with evidence from primary immunodeficiencies and humanized mouse infections that highlight early differentiated and highly proliferative TEMs as being required for EBV specific immune control (45, 77, 83), suggest that further differentiated TRMs with more limited expansion capacity are less protective against EBV infection.

## Methods

### Humanized mouse model

Non-obese diabetic (NOD) severe combined immunodeficiency (scid) γ_c_^null^ (NSG) mice and HLA-A*02:01 transgenic NSG (NSG-A2) mice were maintained in ventilated, specific pathogen–free conditions at the Institute of Experimental Immunology, University of Zurich. Newborn pups were reconstituted with human CD34^+^ hematopoietic progenitor cells (HPCs), derived from human fetal liver tissue (HFL) as previously described (45). For each experiment, animals were reconstituted from a single HFL donor and distributed into different experimental groups with a similar ratio of males and females, as well as similar reconstitution levels of human immune cell populations. No differences in titers or expression of key molecules of interest (i.e. CD69, CD103, cytokines or chemokines) were observed by sex. All animal work strictly followed the animal protocols ZH159/2017 and ZH041/2020, licensed by the veterinary office of the canton of Zurich, Switzerland.

### EBV virus infection and course of experiment

Mice were infected with 10^5^ Raji Green units (RGU) of B95-8 Epstein Barr virus expressing luciferase under the control of the latent EBNA2 locus (45, 46) (Luc-EBV). Luc-EBV producer cells were kindly provided by Prof. Dr. Wolfgang Hammerschmidt (Helmholtz Zentrum, Munich, Germany). As initial histological examination of the nasal sinuses of uninfected animals revealed heterogeneous establishment of NALT structures (Figure S1A-C), we pre-treated animals with i.n.-applied Staphylococcus enterotoxin B (SEB, 62.5ng/µl in PBS), to mimic bacterial colonization of mucosal surfaces driving post-natal NALT development in mice (48, 49). A similar approach (intranasal application of *P*. *acnes* bacteria) has been used to establish of NALT structures in NALT-deficient CXCR5^-/-^ mice (84). Both lymphocyte recruitment (Figure S1B-C) and reproducibility of i.n. infections were significantly improved by SEB pre-treatment (Figure S1G). EBV infection occurred intraperitoneally (i.p.) or intranasally (i.n.) two weeks following SEB pre-treatment, while only i.n. infection led to visible and measurable EBV infection in the NALT (Figure S1H-J). To perform i.n. infections, mice were anesthetized with aerosolized isoflurane and Luc-EBV diluted in PBS was applied in 100µl (for i.p. infection) or 20µl (for i.n. infection) and applied by injection or by slow pipetting in both nostrils, respectively. Mice were monitored regularly for four to six weeks after infection. Intravascular staining was performed by i.v. injection of 6µg Alexa Fluor 700-conjugated CD45 (HI30, Biolegend) diluted in 100µl PBS three to five minutes before euthanasia. In each experimental group, three to six biological replicates were tested.

### In vivo bioluminescence imaging

The progression of EBV infection was monitored longitudinally every week and quantitatively measured by in vivo bioluminescence imaging with the IVIS Spectrum Imaging System (PerkinElmer). Animals were anesthetized by aerosolized isoflurane and injected i.p. with 150mg/kg D-Luciferin (Promega) diluted in PBS 10 minutes before imaging. Mice were placed inside the IVIS imaging box and imaged dorsally and ventrally. Representative images were acquired 10-15 minutes after injection for each mouse throughout each experiment to illustrate viral spread within the host. Images for quantification were captured at various time points before the luminescent signal reached the saturation intensity and analyzed with Living image 4.3.1 software (PerkinElmer). Regions of interest (ROI) were set to include the regions with luminescent signal in mice and photon flux (p/s) of light emitted per second within the ROI was measured as the readout.

### T cell depletion experiments

CD8^+^ T cells were depleted weekly, via i.p. injection beginning at week three post infection, with the monoclonal antibody against human CD8 (clone OKT-8; BioXCell). The initial injection was 75 µg, followed by two injections of 50 µg (week 3) and three injections of 50 µg in week four post infection. Mice with over 10% residual CD3^+^CD4^-^ T cells within lymphocytes in blood and spleen were considered not depleted and excluded from analysis. CD103^+^ cells were targeted by a single i.p. injection at week three with 150ug anti-CD103 (Ber-ACT8, Ultra-LEAF purified, Biolegend).

### Quantification of EBV DNA genome in blood and tissue

Total DNA from whole blood was extracted using NucliSENS easyMag (Biomerieux), while solid organs were processed with DNeasy Blood & Tissue Kit (QIAGEN), according to manufacturer’s instructions. TaqMan (Applied Biosystems) real-time PCR was used to quantify EBV DNA as previously described with modified primers for the BamH1 W fragment (5’-CTTCTCAGTCCAGCGCGTTT-3’ and 5’-CAGTGGTCCCCCTCCCTAGA-3’) and a fluorogenic probe (5’-FAM CGTAAGCCAGACAGCAGCCAATTGTCAG-TAMRA-3’). Viral titer concentration was calculated using a standard curve of the international WHO EBV standard, resulting in titers measured in EBV International Units. All samples were performed in duplicates and measured on either ViiATM 7 Real-Time PCR System (ThermoFisher Scientific) or ABI Prism 7300 Sequence Detector (Applied Biosystems). Samples below the lower limit of quantification (LLOQ) of 173 International Units/ml were defined as negative for EBV DNA.

Normalization of viral loads was performed to adjust for differences in EBV infection levels between experiments with humanized mice that were reconstituted from different hematopoietic progenitor cell donors.

### Humanized mouse sample collection

Peripheral blood cells were obtained from animals by tail vein bleeding or terminal heart puncture. Whole blood was lysed twice with 1xACK lysis buffer for five minutes, followed by washing with PBS. Splenocytes were prepared as described above with one time lysis only. NALT, salivary gland, lung and liver tissues were mechanically disrupted into small pieces and enzymatically digested in 2ml of digestion buffer (1 mg Collagenase D (Roche) and 0.2 mg DNase I (Roche) in 2 ml RPMI or DMEM) at 37°C for 30 minutes with agitation. Dissociated tissues were then passed through a 70 μm cell strainer. NALT was lysed once with 1x ACK lysis buffer for three minutes. Other tissues were partly centrifugated in a discontinuous Ficoll gradient (SG, lung) or Percoll gradient (liver, 40% and 70%, Sigma-Aldrich) for 20 minutes at 1000 rpm using a Sorvall ST 40R centrifuge (Thermo Fisher). Cells aggregating at the interface between gradient layers were harvested and washed twice with PBS. Bone marrow cells were flushed out of the femur by short centrifugation. Cells were washed with PBS and passed through a 70μm cell strainer if necessary. Cells from blood and spleen were counted using the AcT diff Analyzer (Beckman Coulter) to aliquot the optimal number of cells for staining and calculation of the total cell numbers for different experimental purposes. Calculation of total cell numbers for NALT samples was done using Accucheck counting beads (Thermo Fisher) as previously described (85).

### Human sample collection

Human tonsil samples were received from anonymous organ donors in the New York-Presbyterian Hospital; six tonsils from four different individuals were collected immediately after surgery from patients undergoing tonsillectomy for chronic inflammation. All tissues were obtained as part of Institutional Review Board-approved protocols within previous investigations (86). Further tonsil samples were received from tonsillectomies performed at the University Hospital of Zurich. All participants provided informed consent in accordance with the Declaration of Helsinki, and the institutional ethics committee approved all protocols used. Samples were frozen and stored in liquid nitrogen until flow cytometric phenotyping. Samples were rested overnight before stimulation experiments. For comparison of resident vs non-resident populations tonsil samples were divided into CD69^+^ and CD69^-^ subsamples using biotinylated anti-human CD69 antibody (FN40, Biolegend), and separation was performed using anti-Biotin Microbeads (Milenyi, 130-090-485) according to manufacturer’s protocol. Gating during later flow cytometric analyses was chosen according to unstimulated controls from the same samples (Figure S8).

### Flow cytometry, antibody and MHC pentamer labeling

For surface staining, cells were incubated with anti-human FcR-block (Miltenyi), live/dead (Zombie NIR / Aqua, Biolegend) and chemokine-receptor antibodies for 15 minutes at room temperature. Cells were subsequently incubated with surface marker antibodies for 30 minutes at 4°C. For intracellular and intranuclear labeling, surface staining was followed by fixation and permeabilization with the Foxp3/Transcription Factor Staining Buffer Set (eBioscience, 00-5523-00) according to manufacturer’s instructions. Antibodies for intracellular targets were incubated for 1 hour at room temperature. All samples were acquired using DIVA software (BD (Becton, Dickinson and Company)) on LSRFortessa/FACSymphony (BD) instruments, and analysis was performed using FlowJo software (Treestar).

Antibodies (clone, brand, fluorophore) used in this study: CD3 (UCHT1, BD, BUV661), CD4 (SK3, BD, BUV469 / S3.5, Thermo Fisher, PE-Cy5.5), CD8 (SK1, Biolegend, BV650), CD19 (HIB19, Biolegend, PE-Cy5), CD27 (LG.3A10, Biolegend, BV650), CD39 (A1, Biolegend, BV711), CD45 (HI30, BD, BUV395), CD45RA (HI100, Biolegend, APC-Fire750/BV785), CD62L (DREG-56, Biolegend, PE-Cy7 / SK11, BD, BV510), CD69 (FN50, Biolegend, BV421), CD103 (Ber-ACT8, Biolegend, BV711), CD107a (H4A3, BD, FITC), CD127 (IL-7R) (A019D5, Biolegend, Alexa Fluor 700 / PE-Dazzle 594), CD279 (PD-1) (EH12.1, BD, BUV737), CD335 (NKp46) (9E2, Biolegend, BV510), CD56 (B159, BD, APC), Blimp-1 (IC36081R, R&D, Alexa Fluor 647 / 6D3, BD, PE-CF594), CCL5 (VL1, Biolegend, PE), Granzyme B (GB11, Biolegend, PE-CF594 / Alexa Fluor 700), HLA-DR (G46-6, BD, PE-CF594 / BV605), IFN-gamma (4S.B3, BD, BV786), KLRG1 (13F12F2, eBioscience, Alexa Fluor 488), Perforin (dG9, BD, PerCP-Cy5.5), TNF-alpha (Mab11, BD, PE-Cy7), anti-mouse TCR beta (H57-597, Biolegend, BV510 /605)

EBV-pentamers specific for BMLF1 (GLCTLVAML, HLA-A*02:01, PE / APC) and LMP2 (CLGGLLTMV, HLA-A*02:01, APC / PE) were purchased from Pro-immune. Pentamers were added 15 minutes prior to surface antibody labelling during fc-blocking and live/dead staining.

### Human TCR-Vβ repertoire analysis using flow cytometry

Human TCR Vβ repertoire analysis was performed using antibodies against the most frequent TCR Vβ variants (covering about 70% normal human TCR Vβ repertoire, as reported by Beckman Coulter “Beta Mark TCR Vbeta Repertoire Kit” (cat.nr.: IM3497)) and following previously published nomenclature (87)). The antibodies were generously provided by Prof. Dr. Roland Martin (University of Zürich, Zürich. Switzerland). In brief, frozen splenocytes were stained as described above using surface antibodies against following TCR Vβ segments (clone, brand, fluorophore): VB3 (CH92, Beckman Coulter, FITC), VB4 (WJF24, Beckman Coulter, PE), VB5.2 (36213, Beckman Coulter, FITC), VB5.3 (3D11, Beckman Coulter, PE), VB7.1 (ZOE, Beckman Coulter, PE), VB7.2 (REA677, Miltenyi, APC), VB9 (FIN9, Beckman Coulter, PE), VB12 (VER2.32.1.1, Beckman Coulter, PE), VB13.1 (IMMU222, Beckman Coulter, PE), VB13.6 (JU74.33, Beckman Coulter, FITC), VB14 (CAS1.1.3, Beckman Coulter, PE), VB16 (TAMAYA1.2, Beckman Coulter, FITC), VB18 (BA62.6, Beckman Coulter, PE), VB20 (ELL1.4, Beckman Coulter, PE), VB21.3 (IG125, Beckman Coulter, FITC), VB22 (IMMU546, Beckman Coulter, FITC), VB23 (AF23, Beckman Coulter, PE)

After initial analysis, we identified the top six TCR Vβ clones expanded in splenocyte-derived CD8^+^ TEMs in three individuals. These were chosen to stain splenocytes and lymphocytes from the NALT of the same mouse for the following TCR Vβ segments (clone, brand, fluorophore) to identify any overlap of clonotypes: VB1 (BL37.2, Beckman Coulter, PE), VB2 (RE654, Miltenyi, PE-Vio770), VB5.1 (LC4, eBioscience, APC), VB8 (56C5.2, Beckman Coulter, FITC), VB11 (RE559, Miltenyi, PE-Vio770), VB17 (E17.5F3.15.13, Beckman Coulter, FITC)

### Generation of EBV-specific TCR transgenic T cells

EBV-specific T cell receptor (TCR) generation and adoptive T cell transfer experiments were performed as previously described (44, 45). Briefly, for each specificity a total of 200’000 TCR^+^CD3^+^ T cells were transferred intravenously into HPC donor-matched recipient mice and monitored longitudinally during the course of EBV infection.

### Single cell RNA sequencing

Single-cell RNA sequencing of up to 10’000 sorted TEMs was performed using 10X Genomics 3’-kit (v3.1) and Illumina Novaseq S1. Sequencing was performed by the Functional Genomics Center Zurich. Analyses were done with the guidance of NEXUS Zurich and following the OSCA handbook and publication (88). Reads were aligned to a combined reference genome comprising human and EBV reference using the STAR incorporated within the Cellranger software (89). Using R 4.0.5 data were quality controlled and normalized to cell cycle genes and within samples before further analyses (90). With corrected values and Poisson residual’s PCA, TSNE and UMAP calculations were done. Subsequent analyses were performed using BioConductor 3.12 unless stated otherwise (88). Unsupervised clustering was performed using the R-implementation of the Phenograph-algorithm (52) and cellular trajectories were calculated with the TSCAN algorithm (53).

### Ex vivo stimulation experiments

*Ex vivo* isolated and rested CD8^+^ T cells or transduced T cells from splenocytes were incubated with R10 (RPMI 1640 + 10% FBS + 1% penicillin/streptomycin + L-glutamine + 20 U/ml of IL-2; Gibco, Thermo Fisher Scientific) alone or R10 containing PMA/ionomycin for two hours, followed by the addition of Brefeldin A and Monensin and further incubation for an additional three hours. Cells were stained for intracellular cytokines and acquired on a BD FACSymphony. For CD107a labelling, this antibody was added at the very start of coculture. Gating was chosen according to unstimulated controls of the same samples (Figure S8).

### Histology

Whole mouse skulls were fixed using 4% formalin before cutting with a diamond blade. Pieces were decalcified in EDTA at a pH of 9.0 for 20-30 minutes at 100°C and embedded in paraffin. Histological staining was performed by an outside laboratory (Sophistolab AG, Muttenz, Switzerland). In brief, three μm sections were processed on a Leica BOND-MAX or Bond-III automated IHC system. Samples were stained with horse radish peroxidase or alkaline phosphatase-labelled antibodies (clone, manufacturer) for 30 minutes at room temperature: rabbit α–human CD20 (SP32, Cellmarque), mouse α-human CD20 (clone L26, Dako), rabbit α-CD3 (SP7, Diagnostic Biosystem), rabbit α–human CD103 (EPR4166(2), Abcam), rabbit α-mouse Lyve-1 (polyclonal (103-PA50AG), RELIATech GmbH), rat α-mouse/human PNAd (MECA-79, Biolegend). For detection of horse radish peroxidase or alkaline phosphatase, BOND Polymer Refine (DAB) or BOND Polymer Refine Red (Fast Red) were used, respectively (both Leica Biosystems).

Images were acquired with the slide imaging PerkinElmer Vectra3 system. Cell segmentation, total cell count, and phenotype quantification was done using the inform 2.5.1 Tissue Finder Advanced Image Analysis Software from PerkinElmer, as previously described (91). For the final analysis, only cells that the software identified with a confidence >90% were included in the quantification.

### Statistics

Data were analyzed and graphed using Prism Software (v9.5.1 or 8.3, GraphPad). If not described otherwise data sets of two were compared using Mann-Whitney U test and data sets of three were compared using Kruskal Wallis test with Dunett’s multiple comparison. Data within same individuals or samples were compared using paired Wilcoxon signed rank test. For graphing either mean and standard deviation or geometric mean and 95% confidence intervals were displayed.

## Acknowledgements

This research was in part supported by Cancer Research Switzerland (KFS-4962-02-2020), CRPP-ImmunoCure of the University of Zürich, the Sobek Foundation, the Swiss Vaccine Research Institute, the Vontobel Foundation, Roche, Novartis, Viracta, the Swiss MS Society (2021–09) and the Swiss National Science Foundation (310030_204470/1, 310030L_197952/1 and CRSII5_180323). We are grateful to Prof. Hans Strauss of London, UK, for generously providing the plasmids used to generate our TCR transgenic cells. We also thank Prof. Wolfgang Hammerschmidt of München, Germany, for providing the producer cells used to generate luciferase-carrying EBV, as described in this manuscript. We also thank Prof. Roland Martin for providing the human TCR-Vβ specific antibodies. We would also like to thank Mathias Bader (University of Zurich, Information Technology, MELS/SIVIC) for generating the graphical abstract.

## Author contributions

DK, KZ and CM conceived and designed the experiments. DK, KZ, YD, LR, PS, AV and OC performed the experiments. DK, KZ and FL analyzed the data. YD, AM, NC and BC contributed reagents and materials. DK, KZ and CM wrote the manuscript.

