## Supplemental Figures for "Epstein Barr virus infection induces tissue-resident memory T cells in mucosal lymphoid tissues"

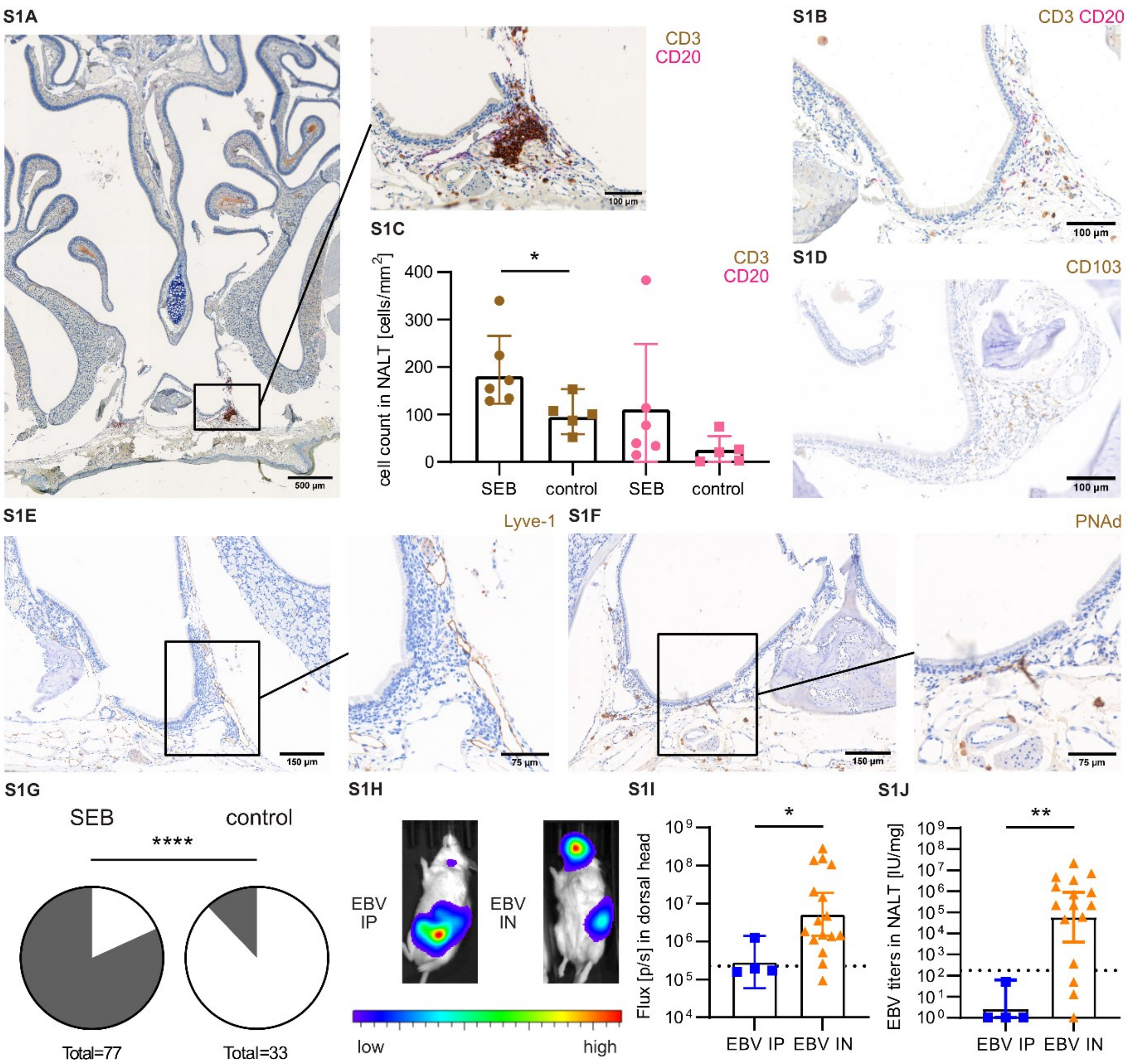


**Supplemental Figure 1: NALT contains B and TRM cells as well as lymphatic vessels/Lyve-1⁺ and PNAd⁺ cells**
(A) Histological slices of Paraffin-embedded PFA-fixed NALT sections stained for CD3 (3,3′-Diaminobenzidine/DAB, brown) as well as CD20 (Fast Red) for SEB-pre-treated and (B) not pre-treated mice. (C) Quantification of $CD3^+$ and $CD20^+$ cells in the NALT area of SEB-pre-treated and not pre-treated (control) mice. (D) Histological slices of Paraffin-embedded PFA-fixed NALT sections stained using DAB for CD103, (E) lymphatic vessel endothelial hyaluronan receptor 1 (Lyve-1) or (F) peripheral node addressin (PNAd). (G) Pie chart quantification of successfully intranasally-infected (gray) and not-infected (white) individual mice with and without SEB pre-treatment (infection status = positive titers in blood at end of experiment). (H) Representative IVIS image analysis from week 6+ after Luc-EBV infection and (I) quantification of a defined head region of interest (ROI size see Figure S2A) of the IVIS images (n=4-16, BKG = 225’000). (J) Quantification of EBV viral loads in International Units (IU) / mg in NALT at time of sacrifice (week 6+ after infection, n=4-16, LOD = 173). (G pooled data of 2-8 independent experiments; ****, $P \leq 0.0001$, Fisher’s exact test; C,I-J pooled data from 1-5 independent experiments; *, $P \leq 0.05$, **, $P \leq 0.01$, Mann-Whitney-U test)

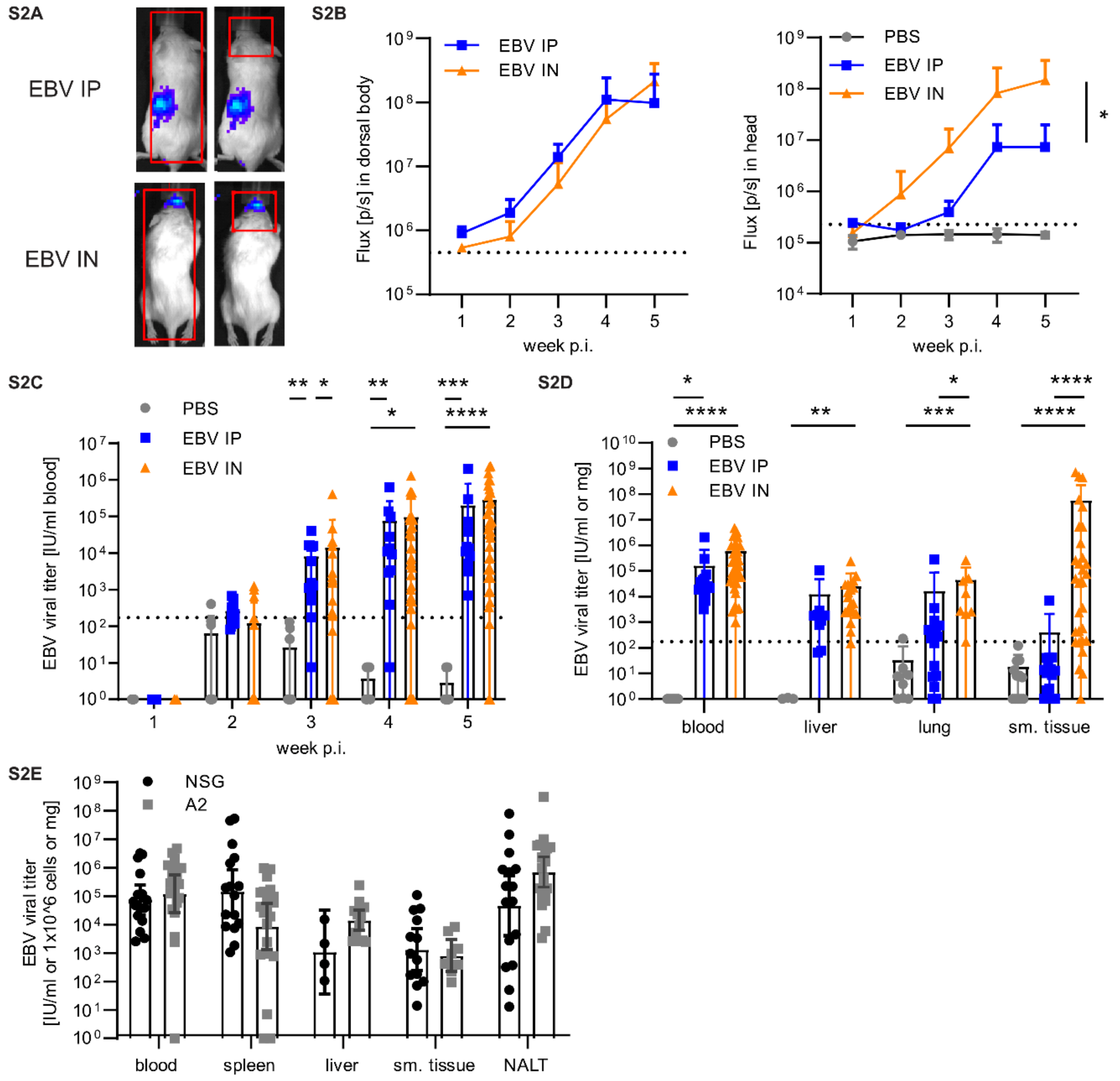


**Supplemental Figure 2: EBV can additionally be detected in NALT and submandibular tissue upon i.n. EBV infection** (A) Representative IVIS image analysis from week 1 after Luc-EBV infection and (B) quantification of a defined region of interest (ROI) of the IVIS images (ROI comprises either the whole body (left, n=9-14, BKG = 450'000) or the head area (right, n=6-24, BKG = 225'000). (C) EBV viral loads in International Units (IU) / ml of animals over time during EBV infection or PBS controls (n=4-31, LOD = 173). (D) Quantification of viral loads in blood, liver, lung or submandibular tissue (n=3-32, LOD = 173). (E) EBV viral loads in IU / ml blood, $1x10^6$ splenocytes or mg tissue at time of sacrifice. (B,C,D pooled data from 2-6 independent experiments; E pooled data of 4 independent experiments for humanized NSG and NSG-A2 (A2); *, P ≤ 0.05 **, P ≤ 0.01; ***, P ≤ 0.001, ****, P ≤ 0.0001, Kruskal-Wallis test including Dunn's multiple comparison test)

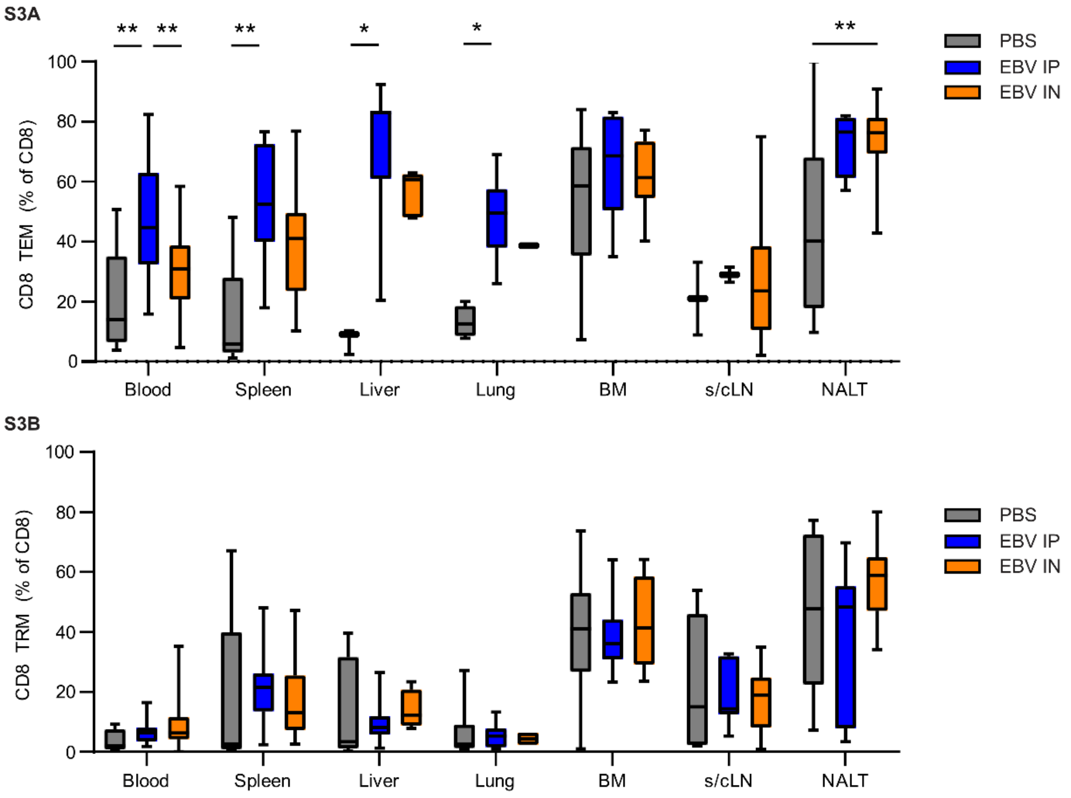


**Supplemental Figure 3: NALT contains B and TRM cells that increase upon intranasal EBV infection**
(A) Frequency of CD8$^+$ TEM populations and (B) CD69$^+$ CD8$^+$ TRM populations in collected organs in PBS (gray), i.p. infected (EBV IP, blue) and i.n. infected (EBV IN, orange) individuals. Pooled data from min. two independent experiments (*, P ≤ 0.05; **, P ≤ 0.01, ***, P ≤ 0.001, Kruskal Wallis test with Dunn's multiple comparison).

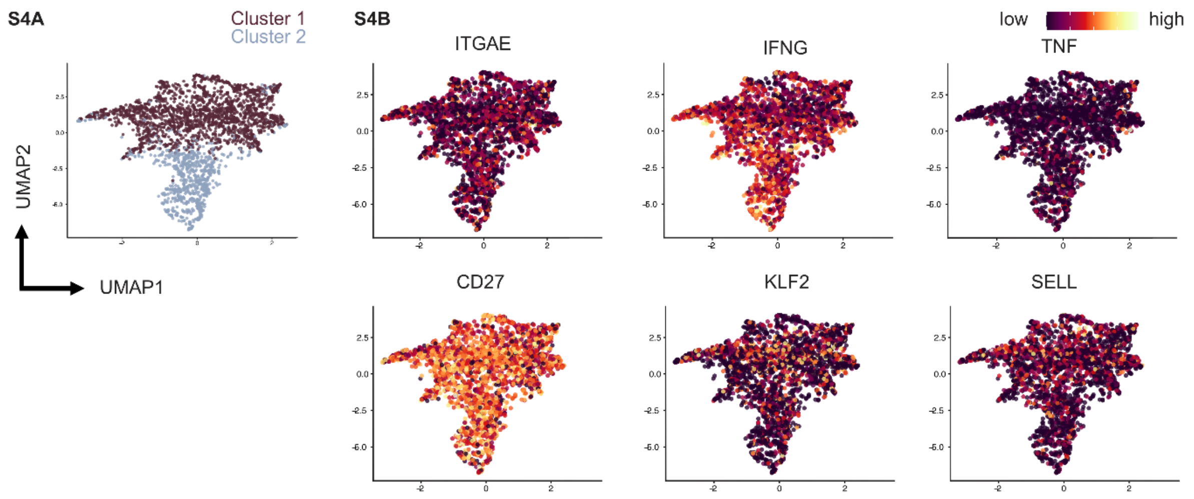


**Supplemental Figure 4: Single cell-RNA-Sequencing of NALT shows heterogenic expression of differentially expressed genes**

A) Integrated UMAP plot of only NALT TEM $CD8^+$ cells showing clustering and (B) relative expression of chosen TRM and circulating T cell marker expression (yellow = high expression, red = intermediate expression, black = low expression).

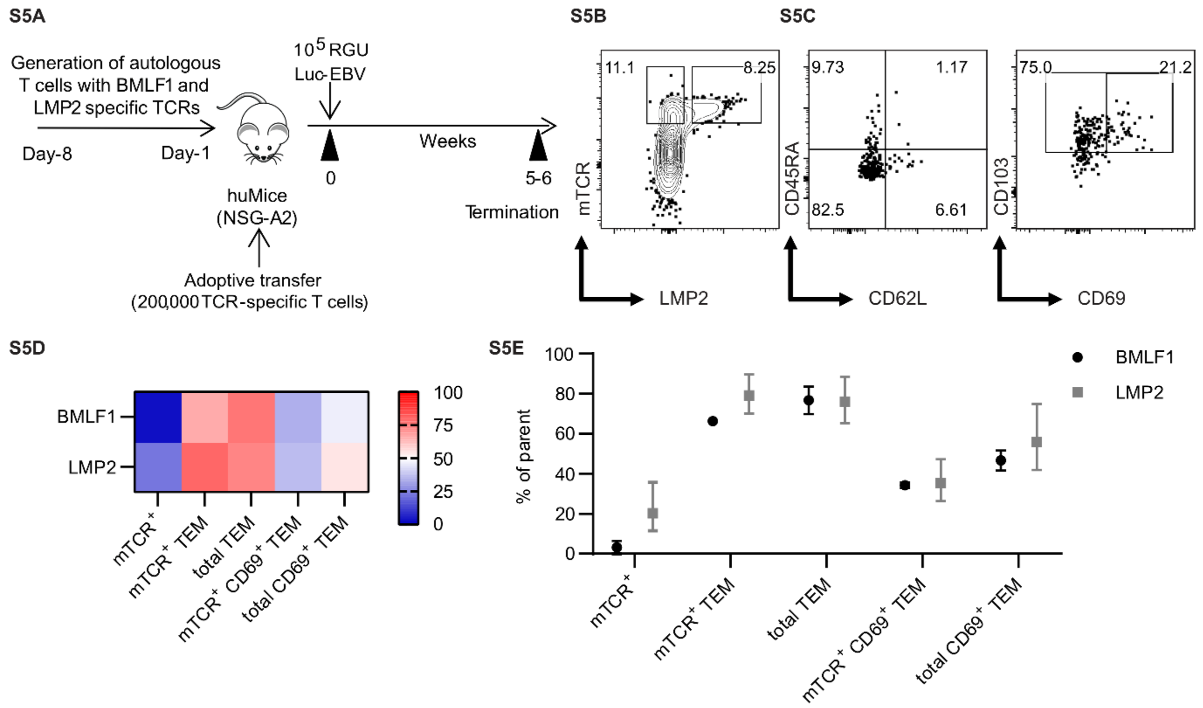


**Supplemental Figure 5: Intranasal EBV infection is capable of inducing EBV-specific T cells with a TRM phenotype**

(A) Experimental layout for EBV-specific T cell experiment. One day before infection with $10^5$ RGU Luc-EBV, huNSG-A2 mice (HLA-A2 transgenic NSG mice reconstituted with HLA-A2$^+$ human hematopoietic progenitor cells) were adoptively transferred with 200’000 autologous T cells expressing either BMLF1- or LMP2-specific TCRs that had been transduced *ex vivo*. (B) Representative flow cytometry plot identifying BMLF1-specific T cells harboring hybrid human/murine transgenic TCR (mTCR) with HLA-A*02:01 plus LMP2 peptide pentamer (LMP2) in the NALT. (C) Representative flow cytometry plots of EBV-specific T cells showing TEM and TRM phenotypes (D). Heatmap and (E) quantification of EBV-specific TCR-T cell populations in the NALT.

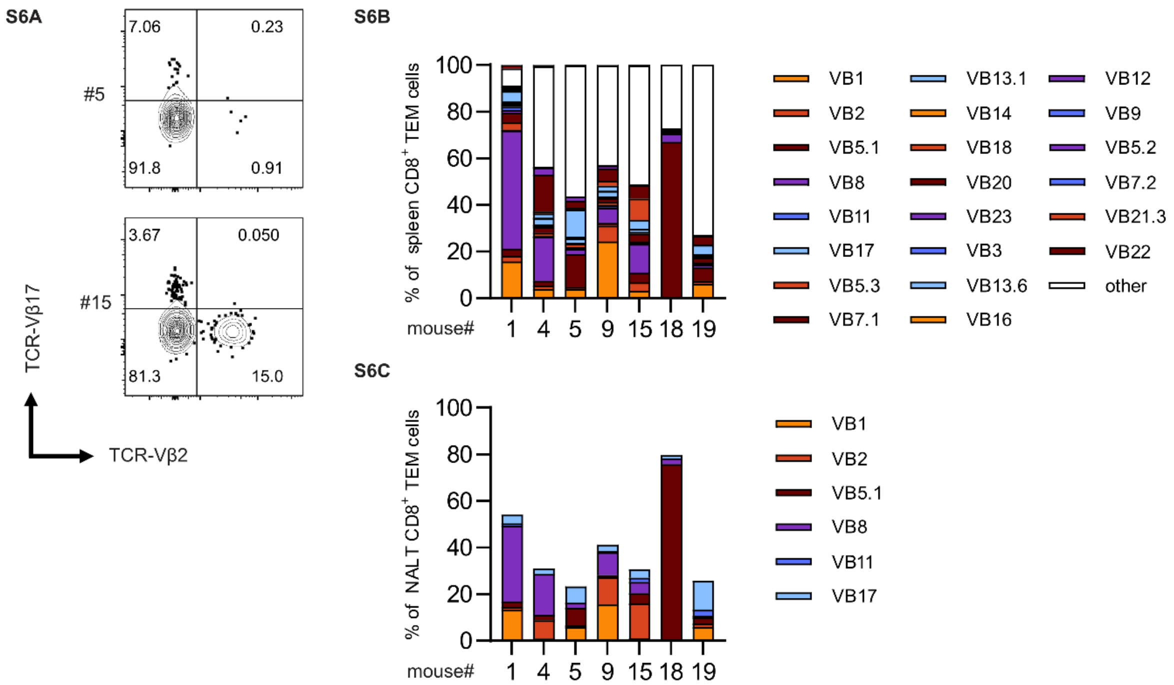


**Supplemental Figure 6: TCR Vβ clonotype analysis shows differential usage of Vβ chains between individuals but shared prominent chains in spleen and NALT TEMs within an individual**
(A) Representative flow cytometry plots of example staining of TCR-Vβ17 and TCR-Vβ2 of two distinct individual humanized NSG mice. (B) Quantification of TCR-Vβ family percentage of spleen $CD8^+$ TEMs. (C) Quantification of the six top used TCR-Vβ clones in NALT $CD8^+$ TEMs of the same individuals.

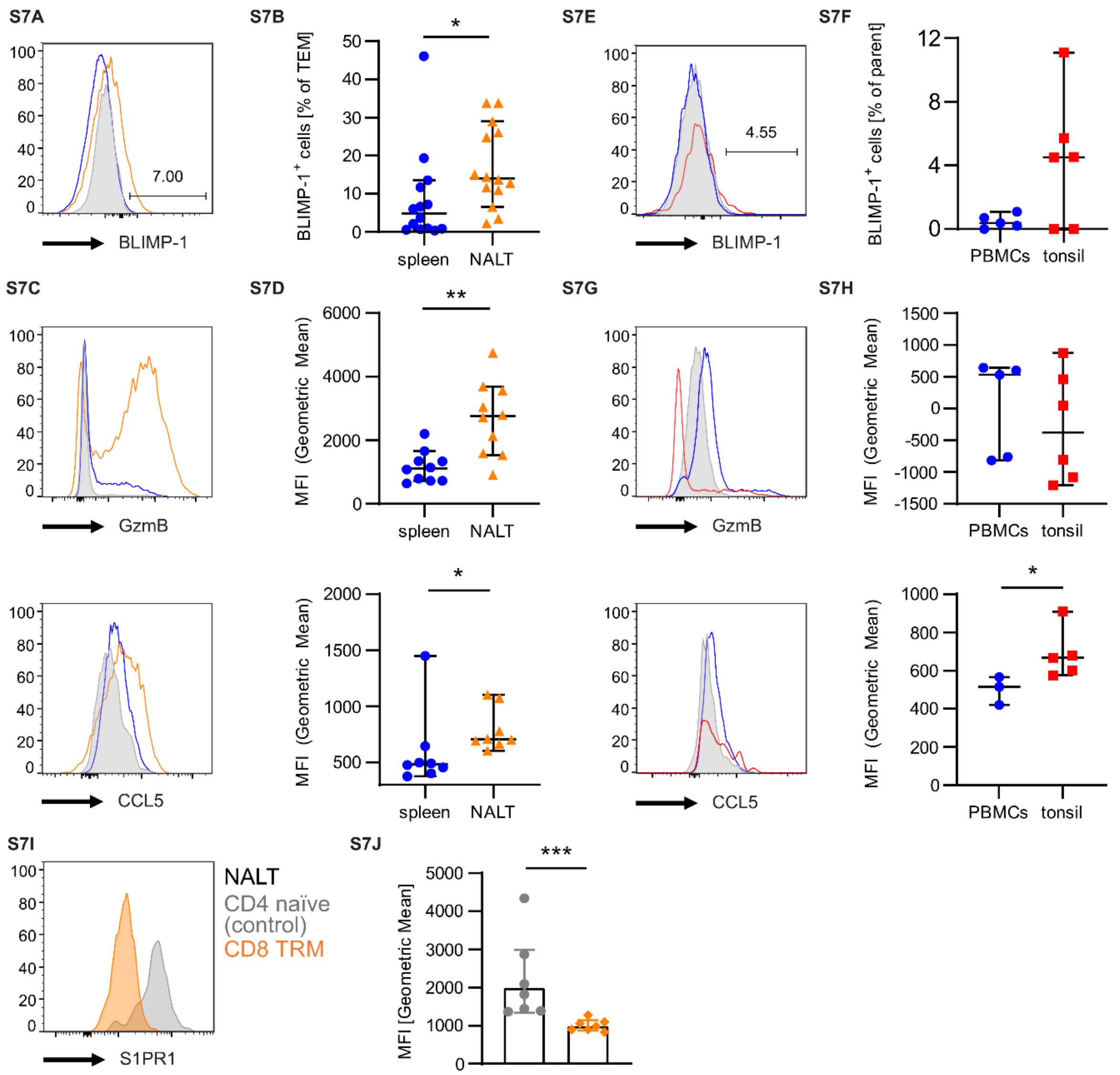


**Supplementary Figure 7: TRM phenotype in NALT and human tonsils**
(A,C) Histograms, (B) quantification of Blimp-1$^+$ TEMs and (D) quantification of geometric mean fluorescence intensity (MFI) of indicated phenotypic markers in TEMs from NALT (orange), TEMs from spleen (blue) and naïve CD8$^+$ T cells from spleen (gray). (E,G) Histograms, (F) quantification of Blimp-1$^+$ TEM and (H) quantification of geometric mean fluorescence intensity (MFI) of indicated phenotypic markers in CD69$^+$CD103$^+$ TRMs from tonsils (red) and TEMs from PBMCs (blue). (I) Histogram and (J) quantification of the MFI of S1PR1 expression in CD8$^+$ TRM (orange) and naïve CD4 T cells (gray) in the NALT. To determine significance, Wilcoxon paired signed rank test (for NALT and spleen) or Mann-Whitney U-test (for tonsil and PBMCs) were performed (*P < .05, **P < .01) and plots represent at least two independent experiments of 7-8 murine or 5-6 human subjects (except data for CCL5 in tonsil/PBMC is from 3-5 human subjects within one experiment and data for S1PR1 is from one experiment).

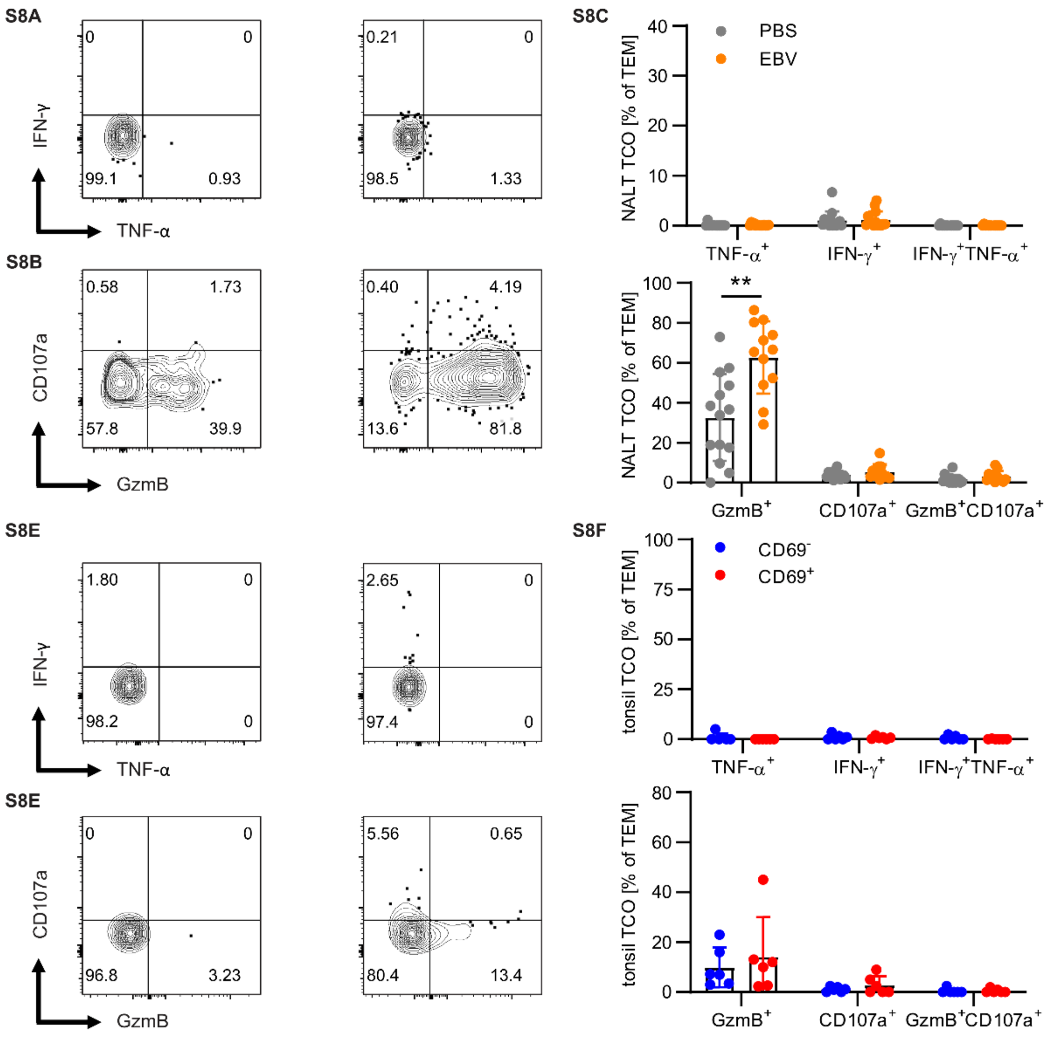


**Supplemental Figure 8: Unstimulated TRMs in NALT and tonsils do not produce cytokines or degranulate *in vitro***
(A) Representative flow cytometry-plots showing IFN-γ and TNF-α or (B) CD107a and Granzyme B (GzmB) expression by $CD8^+$ TEMs in PBS or EBV-infected NALT without stimulation. (C) Quantification in PBS (gray) and EBV-infected (orange) NALT TEMs. (D) Representative flow cytometry-plots showing IFN-γ and TNF-α or (E) CD107a and GzmB expression by $CD69^-$ or $CD69^+$ tonsil $CD8^+$ TEMs. (F) Quantification in $CD69^-$ (blue) and $CD69^+$ (red) tonsil $CD8^+$ TEMs. TCO = T cell only control; data of at least 2 independent experiments (**P < .01, Wilcoxon paired signed rank test)

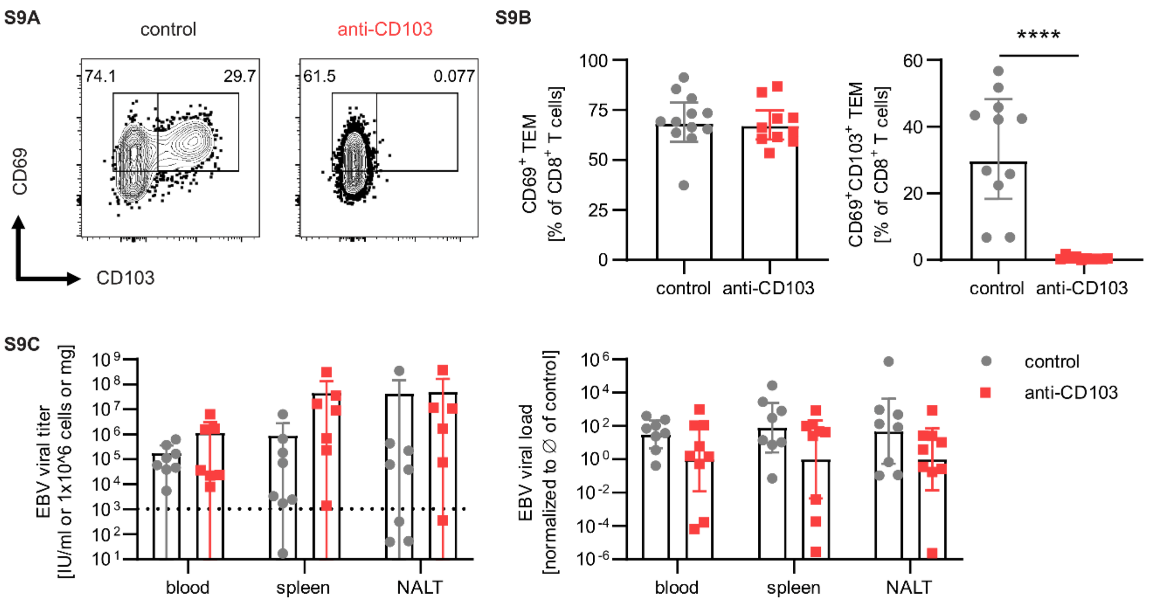


**Supplemental Figure 9: Upon Ber-OCT3 (anti-CD103) treatment, EBV viral loads were unchanged**
(A) Representative plots and (B) quantification of $CD69^{+}$ and $CD103^{+}$ TEM cells in the NALT with or without treatment with Ber-OCT3 (anti-CD103) (C) EBV viral loads in International Units (IU) / ml blood or 1x10^6 splenocytes or mg tissue of NALT (left) or normalized to the mean of the control group per experiment (right). Dotted line is LOD, n=10-12 in two independent experiments (****, $P \leq 0.0001$, Mann-Whitney-U test).
